# Early life exposure to broccoli sprouts confers stronger protection against enterocolitis development in an immunological mouse model of inflammatory bowel disease

**DOI:** 10.1101/2023.01.27.525953

**Authors:** Lola Holcomb, Johanna M. Holman, Molly Hurd, Brigitte Lavoie, Louisa Colucci, Benjamin Hunt, Timothy Hunt, Marissa Kinney, Jahnavi Pathak, Gary M. Mawe, Peter L. Moses, Emma Perry, Allesandra Stratigakis, Tao Zhang, Grace Chen, Suzanne L. Ishaq, Yanyan Li

**Affiliations:** Graduate School of Biomedical Sciences and Engineering, University of Maine, Orono, Maine, USA 04469; School of Food and Agriculture, University of Maine, Orono, Maine, USA 04469; Larner College of Medicine, University of Vermont, Burlington, Vermont, USA 05401; Department of Biology, Husson University, Bangor, Maine, USA 04401; Department of Biology, University of Maine, Orono, Maine, USA 04469; Finch Therapeutics, Somerville, Massachusetts, USA 02143; Electron Microscopy Laboratory, University of Maine, Orono, Maine, USA 04469; School of Pharmacy and Pharmaceutical Sciences, SUNY Binghamton University, Johnson City, New York, USA 13790; Department of Internal Medicine, University of Michigan Medical School, Ann Arbor, Michigan, USA 48109

**Keywords:** Crohn’s Disease, cruciferous vegetables, sulforaphane, glucoraphanin, gut microbiota, dietary bioactives, 16S rDNA, interleukin-10 knockout

## Abstract

Crohn’s Disease (CD) is a presentation of Inflammatory Bowel Disease (IBD) that manifests in childhood and adolescence, and involves chronic and severe enterocolitis, immune and gut microbiome dysregulation, and other complications. Diet and gut-microbiota-produced metabolites are sources of anti-inflammatories which could ameliorate symptoms. However, questions remain on how IBD influences biogeographic patterns of microbial location and function in the gut, how early life transitional gut communities are affected by IBD and diet interventions, and how disruption to biogeography alters disease mediation by diet components or microbial metabolites. Many studies on diet and IBD use a chemically induced ulcerative colitis model, despite the availability of an immune-modulated CD model. Interleukin-10-knockout (IL-10-KO) mice on a C57BL/6 background, beginning at age 4 or 7 weeks, were fed a control diet or one containing 10% (w/w) raw broccoli sprouts, which was high in the sprout-sourced anti-inflammatory sulforaphane. Diets began 7 days prior to, and for 2 weeks after inoculation with *Helicobacter hepaticus,* which triggers Crohn’s-like symptoms in these immune-impaired mice. The broccoli sprout diet increased sulforaphane in plasma; decreased weight stagnation, fecal blood, and diarrhea associated; and increased microbiota richness in the gut, especially in younger mice. Sprout diets resulted in some anatomically specific bacteria in younger mice, and reduced the prevalence and abundance of pathobiont bacteria which trigger inflammation in the IL-10-KO mouse, for example; *Escherichia coli* and *Helicobacter*. Overall, the IL-10-KO mouse model is responsive to a raw broccoli sprout diet and represents an opportunity for more diet-host-microbiome research.

**Importance:** To our knowledge, IL-10-KO mice have not previously been used to investigate the interactions of host, microbiota, and broccoli, broccoli sprouts, or broccoli bioactives in resolving symptoms of CD. We showed that a diet containing 10% raw broccoli sprouts increased the plasma concentration of the anti-inflammatory compound sulforaphane, and protected mice to varying degrees against negative disease symptoms, including weight loss or stagnation, fecal blood, and diarrhea. Younger mice responded more strongly to the diet, further reducing symptoms, as well as increased gut bacterial community richness, increased bacterial community similarity to each other, and more location-specific communities than older mice on the diet intervention. Crohn’s Disease disrupts the lives of patients, and requires people to alter dietary and lifestyle habits to manage symptoms. The current medical treatment is extremely expensive, and a dietary intervention represents an affordable, accessible, and simple strategy to reduce the burden of symptoms.

## Introduction

Inflammatory Bowel Diseases (IBD) affect over 6 million people globally, with more than 25% of cases reported in the United States (1). The varied combination of symptoms, the intensity of their presentation, and their multifactorial origin make symptoms difficult (2) and extremely expensive for patients to manage (3), and detracts from quality of life (4, 5). Crohn’s Disease (CD) is one of the primary immune-disordered presentations of IBD, which typically presents during late adolescence and early adulthood (6), has strongly associated genetic (7) and environmental (8, 9) risk factors, and involves a loss of function of the innate immune system which disrupts host-microbial interactions in the gastrointestinal (GI) tract (10). CD treatments attempt to suppress the immune response to alleviate inflammation as a method of reducing various symptoms, and to return patients to as close to homeostasis as possible. However, some patients (11) and mouse models (12) respond poorly to single-strategy treatments. Diet can play an important, economical, and accessible role in the prevention and/or management of IBD as a source of anti-inflammatory metabolites (13, 14), and broadly for influencing the robustness of the gut microbiome (15–17). However, there are gaps in knowledge about how IBD affects gut microbial ecology, including taxonomic and metabolic biogeography (18–20), how gut microbiota mediate response to disease (21) or diet (22, 23), and how dietary components and microbial metabolites improve symptoms (24, 25).

### Inflammation in CD is severe and disrupts host-microbial interactions

Inflammation in CD is chronic, relapsing, occurs throughout the GI tract, and is complicated to treat because it involves several breakdowns of the innate immune system (10, 26): decreased expression of the MUC1 gene reduces coverage of mucin in the ileum (27) and reduces efficacy of tight junction proteins (28), which can allow microbial translocation from the gut to other tissues (10, 26). Additionally, poor absorption of bile salts in the ileum then causes damage to the colon (29). CD patients mount weak acute inflammation and low neutrophil counts in response to infection (10), and may be unable to clear infections. This results in infiltration of fecal material through the mucosal lining of the gut, dysregulation of the adaptive immune response, and results in chronic inflammation (10, 30). The adaptive immune response in CD primarily involves excessive recruitment of effector T-cells (Th1 and Th17) by inflammatory cytokines (interleukins –12, –18, and –23) which are upregulated in CD lesions (6, 31, 32).

The duration of inflammation, damage to intestinal walls, and other negative outcomes associated with CD increase the risk of developing gastrointestinal cancers, such as colorectal cancer (33). While CD symptoms can develop and present at any age, they are most commonly observed in individuals between the ages of 15 and 30, and approximately 20-30% of cases occur in children and adolescents under the age of 18 (6, 8). Children and adolescents diagnosed with CD encounter distinct challenges, as their gut microbiome is disrupted during critical periods of anatomical, immunological, and microbiological development (34–36). Children experience additional complications such as malnutrition, anemia, impaired growth and development, psychological disorders, bone demineralization, and delayed puberty (37). Dietary treatments for CD could ameliorate age-related aspects of disease (37).

### Diet can be a source of anti-inflammatories to complement medical strategies

Diets which are high in raw, cooked, or fermented cruciferous vegetables can reduce inflammation and cancer risk in the gut of many people (14, 38), in part because these vegetables contain sulfur-containing glucosinolates that can be metabolized to other beneficial compounds (39, 40). Isothiocyanates are a class of bio-active compounds derived from glucosinolates, which can reduce inflammation including in IBD patients (14, 40, 41). Sulforaphane (SFN), the most well-studied isothiocyanate, is produced from the glucosinolate glucoraphanin (GLR), and inhibits the immune factor NF-kB and downregulates multiple inflammatory signaling pathways *ex vivo* and *in vivo* (14, 41–44).

Purified GLR or SFN have been demonstrated to protect against chemical-induced ulcerative colitis in mice (38, 45–47) and some pathogenic bacteria in the gut (48, 49). However, purified SFN is unstable (50, 51), and pure GLR induces extremely variable rates of conversion in humans (40, 51). The conversion of GLR to SFN is achieved by plant-sourced enzymes when raw vegetables are chewed or chopped, and by microbially sourced enzymes in the gut especially if vegetables are cooked to heat-inactivate plant enzymes (38, 46, 49). GLR is concentrated in broccoli and especially immature sprouts, and a whole-food strategy using broccoli sprouts increases gut microbial richness, increases sulforaphane in colon tissues and plasma, and reduces inflammation in chemically induced models of colitis (38, 45, 46). Previous diet research used mouse models lacking the complex immune-dysfunction component of CD (52, 53). Research must elucidate how the immune system of CD patients will react to diet-induced microbiota changes (24, 54), as CD patients have historically been advised to avoid cruciferous vegetables (54) and sulfur-rich diets (55).

### CD disrupts microbial community development from adolescence to early adulthood

The composition of the gut microbiota changes from transitional microbial communities in childhood to a more stable community in adulthood (16), with distinct variations in the composition of gut microbiota between people who are healthy or have CD. For example, pediatric patients exhibit decreased bacterial α-diversity and β-diversity compared to healthy children (34–36)), and while no specific gut microbiota alterations were consistently reported, a gain in *Enterococcus* and a significant decrease in *Anaerostipes, Blautia, Coprococcus, Faecalibacterium, Roseburia, Ruminococcus*, and *Lachnospira* have been noted (34–36). There are also variations across different age groups with CD, from infants through elderly patients (56–58). Older patients tend to exhibit a higher prevalence of colitis, whereas younger patients are more likely to present with ileal disease (59), which may result in age-specific and location-specific changes to gut microbiota. A more thorough understanding of microbial location in the gut (biogeography) may provide more insight into the interactions between host and microbiota in the development of CD.

### Diet studies using immunological models of IBD are lacking

The interleukin 10 (-/-) knockout (IL-10-KO) mouse is commonly used as a non-chemical, genetic model well-suited to studying the immune factors, inflammation, and microbiota of CD (60, 61). Interleukins (IL) are signaling proteins that regulate inflammatory and immune responses and mediate host-microbial tolerance (62). IL-10 also stimulates the growth and differentiation of numerous cell types, suppresses macrophage activation, inhibits inflammatory cytokine production, displays multiple mechanisms of control of Th1 cells (63), and influences the innate immune response to microorganisms without which there is breakdown of host-microbial relations in IBD (64). IL-10-KO are raised in pathogen-free conditions and develop chronic enterocolitis upon exposure to microorganisms which act commensally in immune-competent mice, and their response resembles the transmural inflammation of CD, complete with the formation of granulomas, crypt abscess, mucosal hyperplasia, as well as aberrant immune cell response (65, 66).

To our knowledge, IL-10-KO mice have not previously been used to investigate the interactions of host, microbiota, and broccoli, broccoli sprouts, or broccoli bioactives in reducing inflammation, modifying the immune response, and supporting GI tract microbial systems in CD patients. The objective of this study was to evaluate IL-10-KO mice as a model for studying the broccoli sprout bioactives. We hypothesized that a medium-high concentration (10% w/w) of raw broccoli sprouts in the diet, which contains both the glucoraphanin (GLR) precursor and the anti-inflammatory byproduct SFN, would protect mice from the effects of inflammation triggered by microbial conventionalization even in a host with disrupted responses to commensal bacteria. We hypothesized that sprouts would alter the gut microbiota and in location-specific patterns, and increase the abundance of potentially beneficial taxa while reducing the abundance of putative pathogens.

It is standard to conventionalize IL-10-KO at approximately 8 weeks of age (67), to mimic adolescence and typical timing for the onset of symptoms. In conducting two replications of a diet trial, logical constraints during the pandemic resulted in the use of mice beginning the diet at 4-weeks (∼1 week after weaning (68)) and 7-weeks of age, and conventionalized at 5 and 8 weeks, respectively. We observed a previously unreported effect of young age on the effectiveness of the response to broccoli sprouts in the diet in resolving the symptoms of enterocolitis, and added an *a posteriori* hypothesis that younger mice with transitional gut microbial communities might be more amenable to changes from this diet (68, 69).

## Results

### Raw broccoli sprouts alleviated disease characteristics of immune-modulated enterocolitis

The experimental design was structured as a prevention paradigm (Figure 1) in which specific-pathogen-free, homozygous IL-10-KO mice were given *ad libitum* either a standard chow diet (5LOD irradiated) or the treatment diet consisting of 10% (w/w) raw broccoli sprouts, which were balanced for micronutrients and fiber. Mice were acclimated to diets for 7 days, continuing for a further 16 days during the induction of colitis via microbial colonization and symptom onset. Raw broccoli sprouts contain concentrated amounts of GLR, some of which is metabolized to SFN by mastication, or in this case, diet preparation, when the broccoli enzyme myrosinase is released from tissues (70). The control diet contained no GLR or SFN, and the broccoli sprout diet contained on average 4 μg of GLR and 85 μg of SFN per gram of diet sampled.

**Figure 1.**
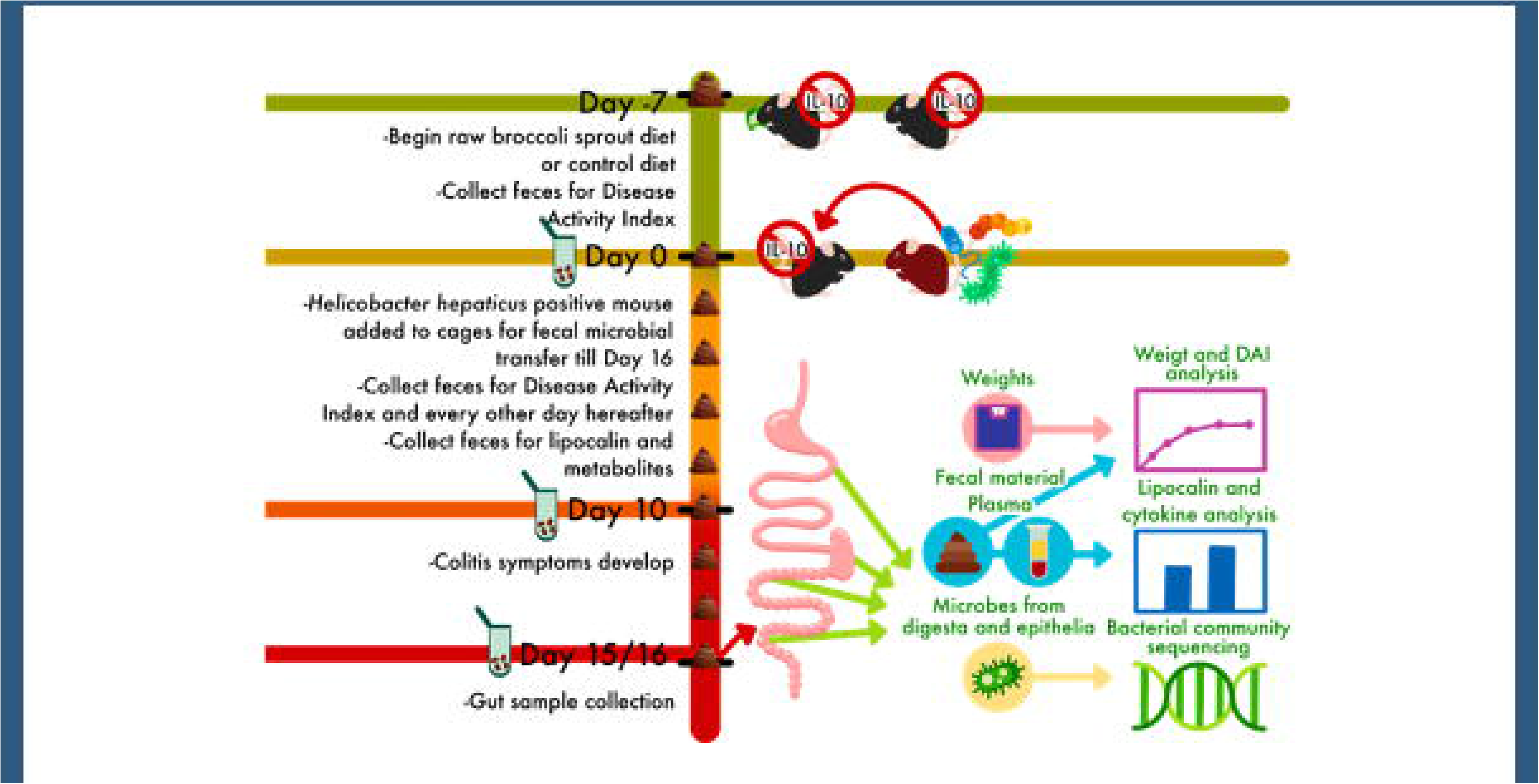
**Experimental design for a dietary prevention model of Crohn’s Disease in Interleukin-10 knockout mice beginning at age 4 or 7 weeks**. Mice were raised in a barrier facility and fed a control or 10% raw broccoli sprout diet for 7 days, then for an additional 16 days as they were moved to a conventional room and co-housed with a *Helicobacter hepaticus* positive mouse to induce immune-mediated enterocolitis.

Trial 1 included 9 mice (n = 5 in the treatment group and 4 in the control group) starting the diet at 4 weeks of age, and being exposed to diverse microbiota at 5 weeks of age. Trial 2 included 11 mice (n = 5 in the treatment group and 6 in the control group) starting the diet at 7 weeks of age, and being exposed at 8 weeks which is the standard age for inducing symptoms in IL-10-KO microbiota trials. The younger mice consuming the broccoli sprout diet continued to gain weight once enterocolitis was induced compared to their baseline weight, and the younger mice fed the control diet plateaued around 120% of body weight (Figure 2A, ANOVA p < 0.05). The older mice had a similar weight stagnation during enterocolitis regardless of diet (Figure 2B).

**Figure 2.**
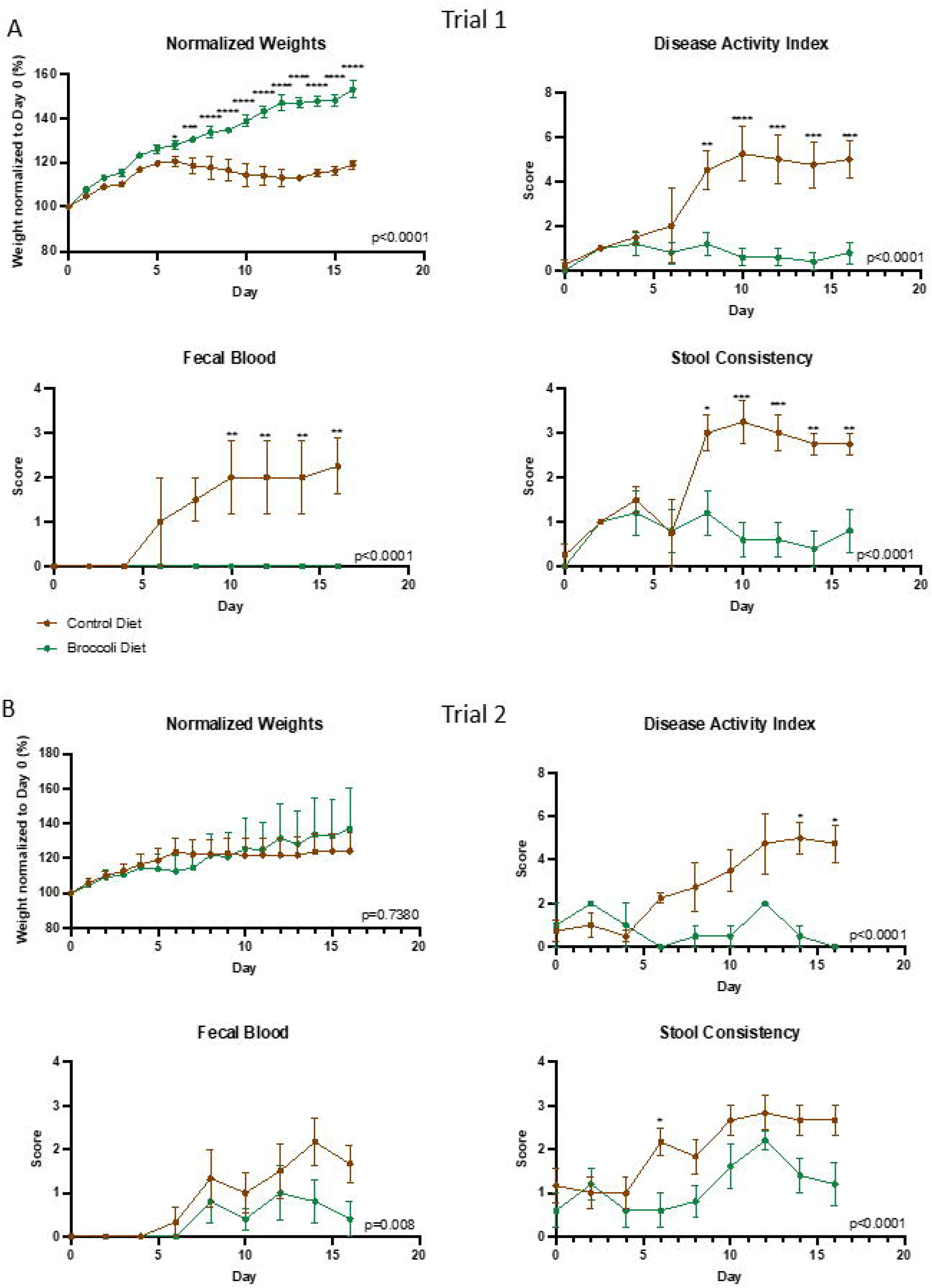
A 10% raw broccoli sprout diet significantly attenuated the development of colitis in IL-10-KO mice that were A) 4 – 6 weeks old during trial or B) 7 – 9 weeks old during the trial. Brown lines indicate control diet (n = 4, trial 1; n = 6 trial 2) and green lines indicate sprout diet (n = 5, trial 1; n = 5, trial 2). All mice were in a growth phase, so body weights and time scale were normalized to the day mice were exposed to *H. hepaticus*-positive mice, set at 100% starting weight and Day 0, respectively. DAI scores are calculated by weight loss intensity score, fecal blood, and fecal consistency. Overall model significance is included on the chart, and single timepoint comparisons significance was designated as p<0.05; **p<0.01; ***p<0.001; ****p<0.0001 by two-way ANOVA, and post multiple comparisons by Sidak.

Consumption of the broccoli diet significantly attenuated the development of enterocolitis in IL-10-KO mice regardless of age group, as indicated by lower Disease Activity Index (DAI) scores (Figure 2, ANOVA p < 0.05). DAI scores were calculated based on three characteristics: fecal blood (presence/absence), fecal blood severity, and fecal consistency (firm/loose/diarrhea). The higher the DAI score, the more severe the disease was.

Lipocalin (LCN2), a neutrophil protein that binds bacterial siderophores, serves as a biomarker for intestinal inflammation (71). Analysis of fecal samples from both age groups on the day of microbial exposure for conventionalization (D0), appearance of symptoms (D10), and end of the trial (D15/16) revealed a markedly lower LCN2 concentration in the group of mice fed the 10% raw broccoli sprouts diet compared to the control diet at both D10 and D15/16 (Figure 3A, ANOVA p < 0.05). This was confirmed by serum lipocalin on D15/16, which showed a marked decrease in LCN2 concentration in the group of mice fed the 10% raw broccoli sprouts diet compared to the control diet (Figure 3B, ANOVA p < 0.05). Similarly on D15/16, pro-inflammatory cytokines IL-1β, IL-6, and TNF-α were significantly reduced in the plasma of mice consuming broccoli sprouts (Figure 3C, ANOVA p < 0.05). CCL4 (known as macrophage inflammatory protein 1β (MIP1β)) which regulates the migration and activation of immune cells during inflammatory processes was not different in younger mice and not evaluated in older mice.

**Figure 3.**
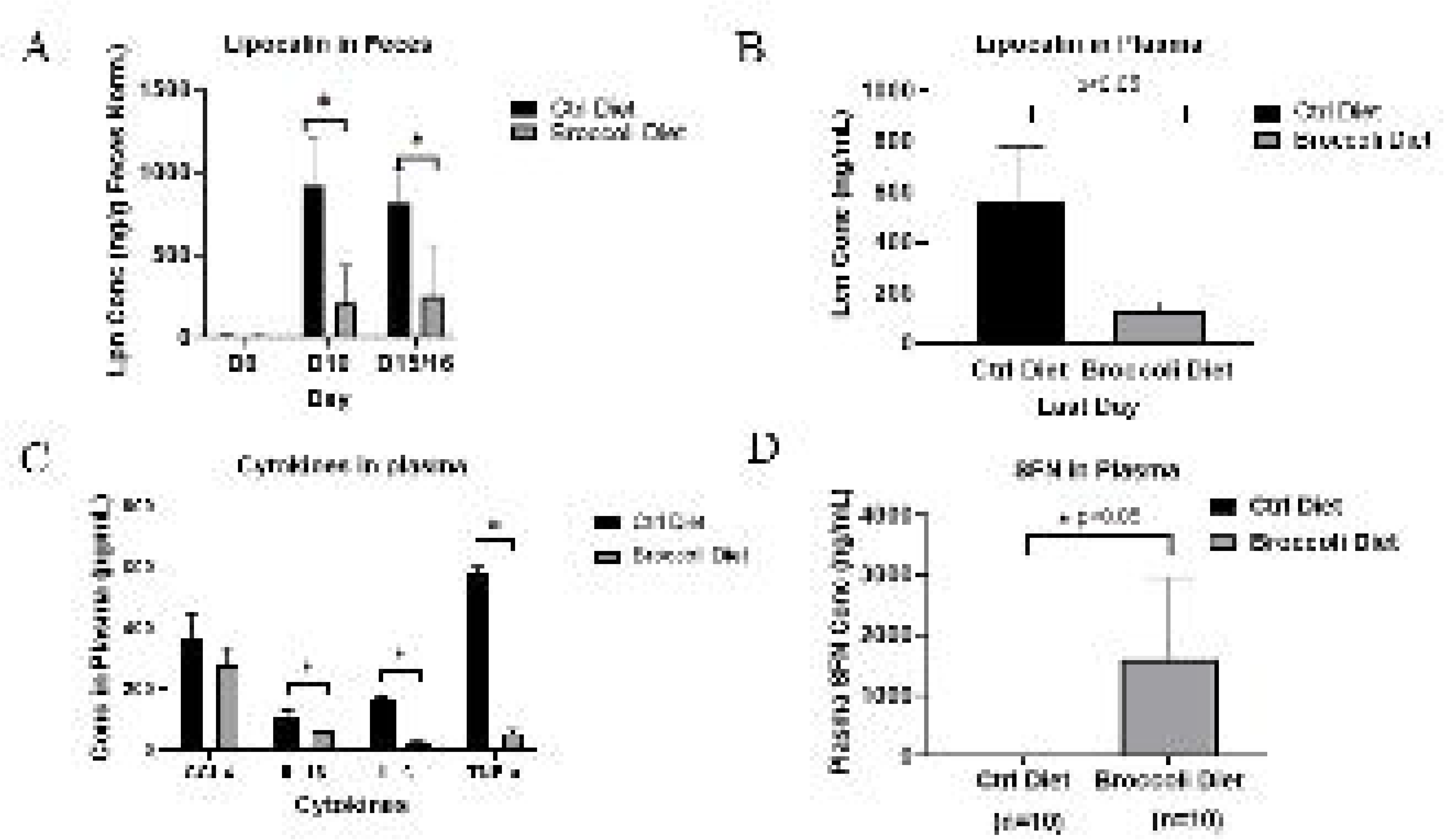
Levels of (A) fecal lipocalin during the diet trial and plasma (B) lipocalin, (C) cytokines, and (D) sulforaphane on the last day, in IL-10-KO mice with enterocolitis during consuming 10% (w/w) raw broccoli sprouts or control diets. Stool samples were collected from mice (n=10/group across both trials) on Days 0 (conventionalization), 10 (onset of symptoms), and 15/16 of experiment for Lipocalin-2 (Lpn/Lcn) concentration determination. Plasma was collected on Days 15/16 for lipocalin, cytokines, and sulforaphane concentration. Data were similar across both trials and combined. For fecal lipocalin, data was normalized by the weight of feces, and significance set as * p<0.05.

Tissue from the ileum, proximal colon, and distal colon were collected for histological scoring (0 = no signs, to 2 = significant signs) of epithelial damage, architectural changes, infiltration of mononuclear cells into the lamina propria, infiltration of polymorphonuclear cells in the lamina propria or into the epithelium, as well as abscesses, ulcers, erosion, and branched crypts which were scored together (Figure S1). Diet was not a significant factor for most scoring criteria, even when data were subset by trial/age, or subset by organ. The exception to this was infiltration of mononuclear cells into the lamina propria, which was significantly lower (linear regression model (lm), p = 0.02) in the younger broccoli sprout mice compared to control mice in trial 1.

SFN was absorbed from the gut by the mice consuming broccoli sprouts in both age groups, and was found in high concentrations in their plasma, while control mice exhibited no circulating SFN (Figure 3D, ANOVA p < 0.05).

### Age significantly influenced bacterial community responsiveness to raw broccoli sprouts

Of the three experimental factors analyzed in this study, we report that age is the most significant factor (relative to diet and anatomical location) in driving the richness of gut bacterial communities, as well as their similarity to each other. The younger trial 1 mice that started consuming the raw broccoli sprouts at 4 weeks of age had greater observed bacterial richness by 6 weeks of age than their counterparts fed a control diet (Figure 4A, permANOVA p < 0.05). However, mice in trial 2, starting at 7 weeks, showed no significant difference in bacterial richness compared to their control group by age 9 weeks (Figure 4B, permANOVA p > 0.05). When comparing bacterial communities within each age group and within each of the four anatomical locations studied, the broccoli sprout diet increased bacterial richness in the cecum and the proximal colon of the younger mice, compared to the younger controls (Figure 4B, Wilcox tests p < 0.05). There was no difference between bacterial richness in any gut location in the older mice consuming broccoli sprouts versus the control diet.

**Figure 4.**
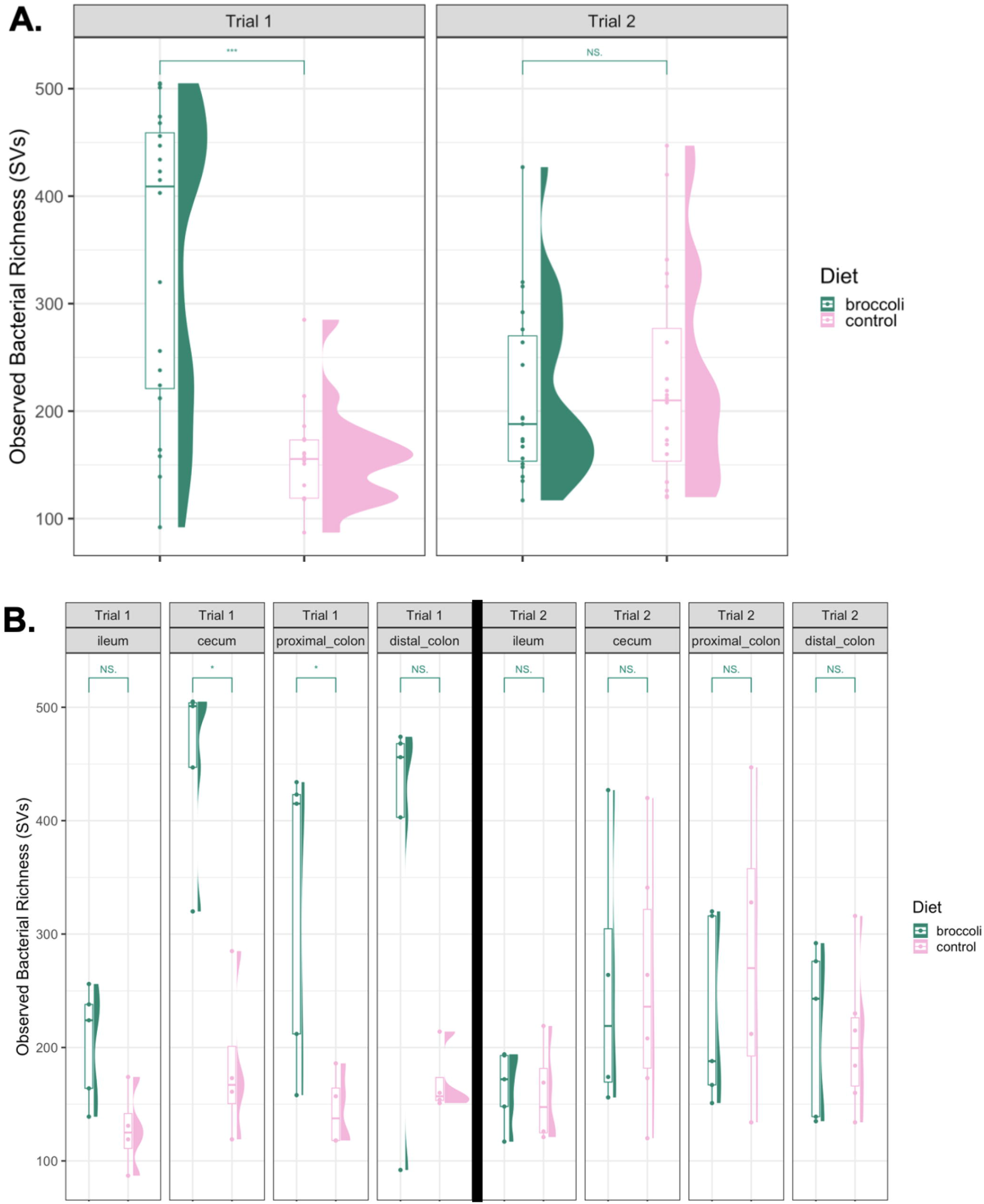
Observed bacterial richness within the gastrointestinal tracts IL-10-KO mice fed control diets or raw broccoli sprout diets from 4-6 weeks of age (Trial 1) and 7-9 weeks of age (Trial 2). Panel A shows comparison of broccoli-fed and control mice in Trial 1 and Trial 2 mice across the whole gastrointestinal tract (***p<0.001); Panel B shows comparisons of broccoli-fed and control mice in which were 4 – 6 weeks old for the duration of the first trial and 7 – 9 weeks old in the second trial, in each anatomical location scraping site (*p<0.05). Graphics made using phyloseq and ggplot2 packages in R. Significance added by Wilcox tests from the ggsignif package in R.

When comparing bacterial community similarity between all samples (beta diversity), age was again the strongest explanatory factor when comparing bacterial taxa presence/absence (unweighted, Figures S2, S3) and bacterial taxa presence and abundance (weighted, Figures 5, 6) metrics. Within the younger trial 1 group, bacterial communities clustered separately by diet (Figure 5; Table 1, permANOVA p < 0.001). This was, in part, due to significant taxonomic dissimilarity between the two diet groups in the cecum and the distal colon (Figure 6, Table 1), while differences between diet groups within the proximal colon were approaching significance. Within the older trial 2 mice, diet was also the most significant driver of bacterial community similarity, but to a lesser extent than in the younger mice (Figure 5; Table 1, permANOVA: lower F values, p < 0.03). Overall, anatomical location was a significant factor in bacterial community similarity in older mice, however; within each of the four anatomical locations studied, there was no significant effect between the diets (p > 0.05), indicating smaller-scale changes across the GI tract communities.

**Figure 5.**
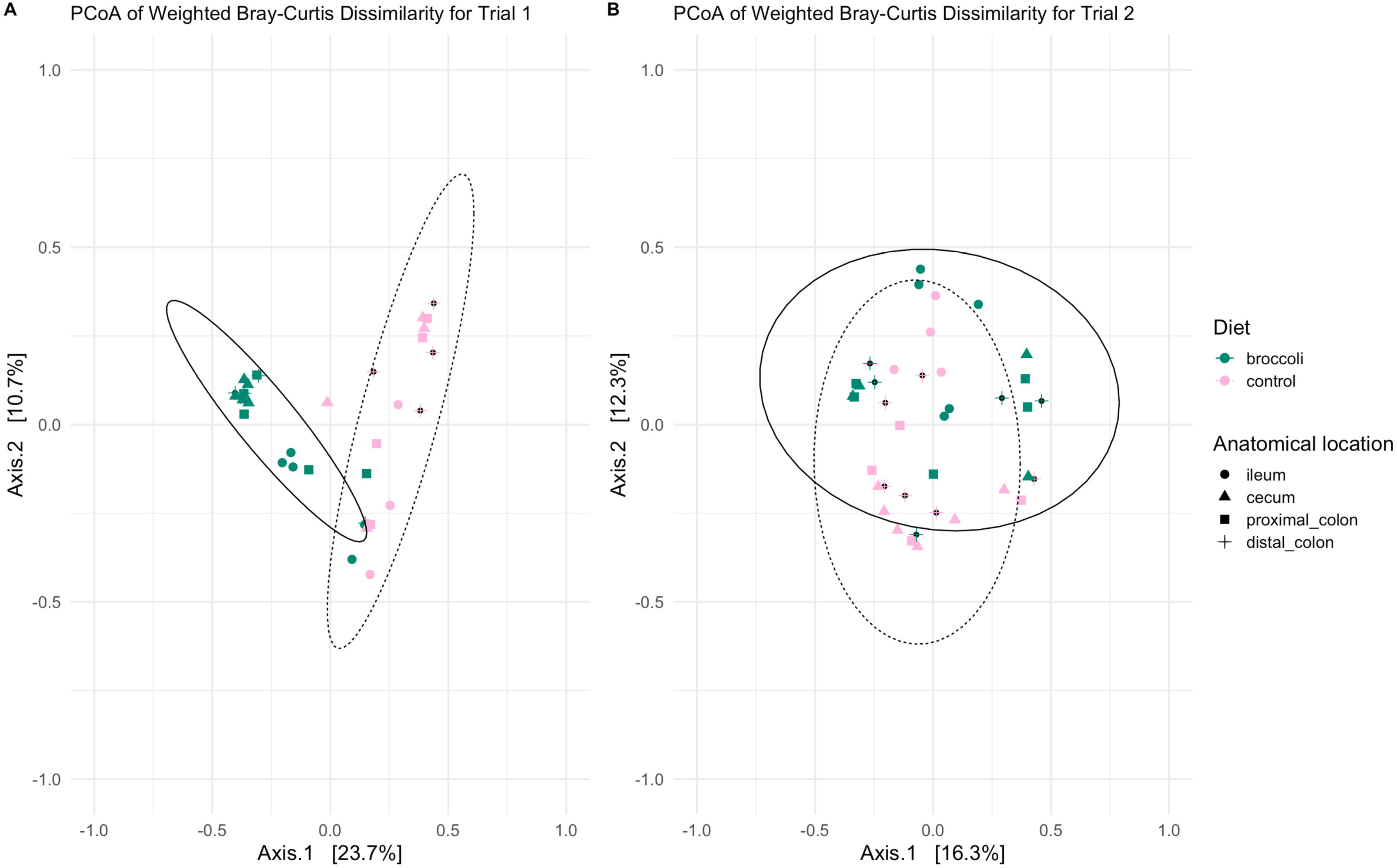
Principal coordinates analysis of bacterial community similarity within the gastrointestinal tracts of 4– or 7-week-old IL-10-KO mice fed control diets or broccoli sprout diets. Calculations using weighted Bray-Curtis dissimilarity show differences in the taxonomic structure of trial 1 mice in Panel A, and trial 2 mice in Panel B. Graphic made using phyloseq, and vegan packages in R.

**Figure 6.**
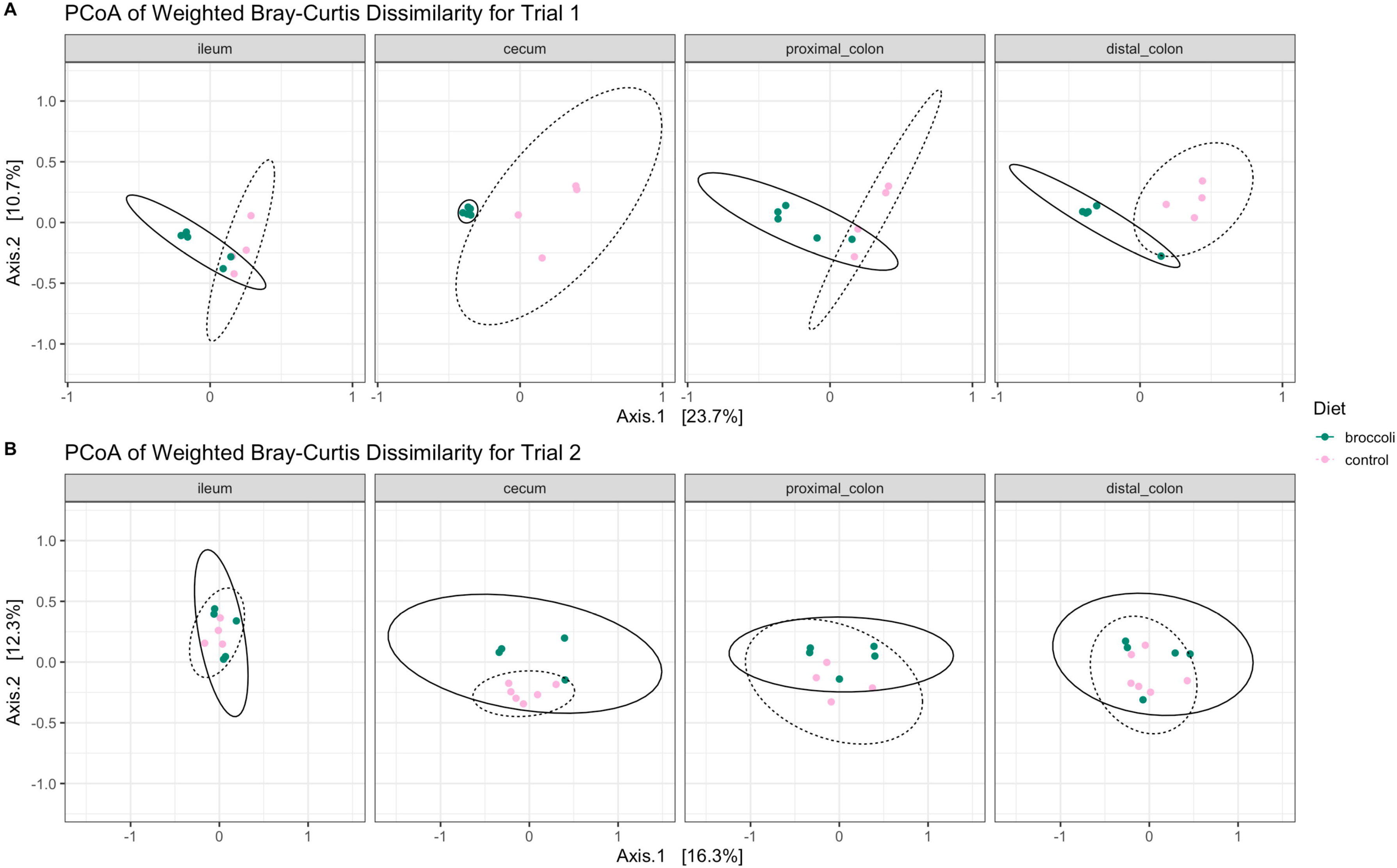
Bacterial community dissimilarity within a specific anatomical location along the gastrointestinal tract of IL-10-KO mice fed control diets or broccoli sprout diets beginning at 4 or 7 weeks of age. Calculations using weighted Bray-Curtis dissimilarity show differences in the taxonomic structure of (A) trial 1 mice sampled at age 6 weeks, and (B) trial 2 mice sampled at age 9 weeks, after each had been consuming diets for 3 weeks. Principal coordinates analysis graphic made using phyloseq, and vegan packages in R.

**Table 1.** Statistical comparison of principal coordinates analysis of bacterial community similarity within the gastrointestinal tracts of 4– or 7-week-old IL-10-KO mice fed control diets or broccoli sprout diets. Comparisons were done using dietary treatment and anatomical locations as factors, and subsetting by trial, as well as by anatomical location, as noted. Unweighted Jaccard Similarity and weighted Bray-Curtis metrics were used to calculate beta diversity. Number of permutations = 9999.

| <u>Permanova Tests of Dissimilarity</u> | Df | Test type | Trial 1, aged 4-6 weeks |  | Trial 2, aged 7-9 weeks |  |
| --- | --- | --- | --- | --- | --- | --- |
|  |  |  | F | P-value | F | P-value |
| Diet | 1 | Jac | 2.7779 | 0.0001*** | 1.1898 | 0.0258* |
|  |  | BC | 7.5867 | 0.0001*** | 2.4498 | 0.0022** |
| Anatomical Location | 3 | Jac | 1.2027 | 0.0291* | 1.0955 | 0.0359* |
|  |  | BC | 1.5991 | 0.0205* | 1.4958 | 0.0139* |
| Diet: Anatomical Location | 3 | Jac | 1.0704 | 0.1780 | 0.9559 | 0.8477 |
|  |  | BC | 1.0482 | 0.3504 | 0.8272 | 0.8300 |
| Diet effect within ileum | 1 | Jac | 1.1319 | 0.1242 | 1.0154 | 0.3133 |
|  |  | BC | 1.4445 | 0.1220 | 1.3366 | 0.1699 |
| Diet effect within cecum | 1 | Jac | 1.9352 | 0.0074** | 1.0393 | 0.2663 |
|  |  | BC | 4.4036 | 0.0081** | 1.6667 | 0.0818 |
| Diet effect within proximal colon | 1 | Jac | 1.3499 | 0.0554 | 0.9949 | 0.3998 |
|  |  | BC | 2.0148 | 0.0247* | 0.8776 | 0.6080 |
| Diet effect within distal colon | 1 | Jac | 1.7018 | 0.0261* | 1.0058 | 0.4183 |
|  |  | BC | 4.1923 | 0.0158* | 0.9797 | 0.4679 |

Significance codes: $p \leq 0.001$ \*\*\* | $p \leq 0.01$ \*\* | $p \leq 0.05$ \*
Jaccard dissimilarity = "Jac" | Bray-Curtis dissimilarity = "BC"

The bacterial taxa which drove the differences between age, diet, and anatomical treatment were primarily different sequence variants (SVs) identified to the Muribaculaceae and Lachnospiraceae families, common mouse commensals, abundant in younger broccoli sprout fed mice (Figure S4). The lactic acid-fermenting genus *Limosilactobacillus* was found in older mice, and was more abundant in older mice consuming broccoli sprouts (Figure S4). *Helicobacter typhlonius*, which is pathogenic in immunocompromised mice, was abundant in control mice of both ages, and in older mice consuming sprouts, while *Escherichia* was found in many locations in the control mice but only a few older mice consuming sprouts (Figure S4).

Bacterial taxa which were abundant in more than 70% of samples in the groups subset by trial/age and diet were considered part of the “core” community. SVs in the Muribaculaceae family made up much of the core microbiota specific to each group, as well as several SVs identified to Lachnospiraceae, *Bacteroidetes*, *Helicobacter*, and others (Figure S5). *Helicobacter* was more prevalent and abundant in control mice of both ages.

The Source Tracker algorithm was used to determine if the cecum could be the ‘source’ for bacterial population ‘sinks’ in the colon, as a proxy for the mouse model’s applicability to the human gut anatomical features and microbial communities. A total of 38 SVs were identified as possibly sourced from the cecum (Figure S6). Common mouse commensals, but not the putative GSL-converting bacteria in the broccoli sprout fed gut samples, were among those taxa identified as sourced in the cecum and sinking in the proximal or distal colon.

### Raw broccoli sprout diet reduced the abundance of putative pathogens

SFN is bacteriostatic against common gut pathogens, including *Escherichia coli*, *Klebsiella pneumonia*, *Staphylococcus aureus*, *S. epidermidis*, *Enterococcus faecalis*, *Bacillus cereus* as well as non-pathogenic *Bacillus,* and *Helicobacter* spp., which can be commensal in immune-competent mice (48, 72). Using that previous literature, we quantified differences in abundance in putative pathogens, which were higher in control groups, across more locations in the gut, and in higher abundance than the broccoli sprout groups (Figure 7). This trend was more pronounced in older control mice which exhibited an abundance of *Helicobacter* SVs through the cecum, proximal colon, and distal colon, as compared to the younger controls (Figure 7, ANOVA p=0.147). Within the broccoli sprout groups, the older mice also appeared to have a greater abundance of *Helicobacter* SVs compared to their younger counterparts in the distal colon and though it was not statistically significant (Figure 7, ANOVA p=0.657).

**Figure 7.**
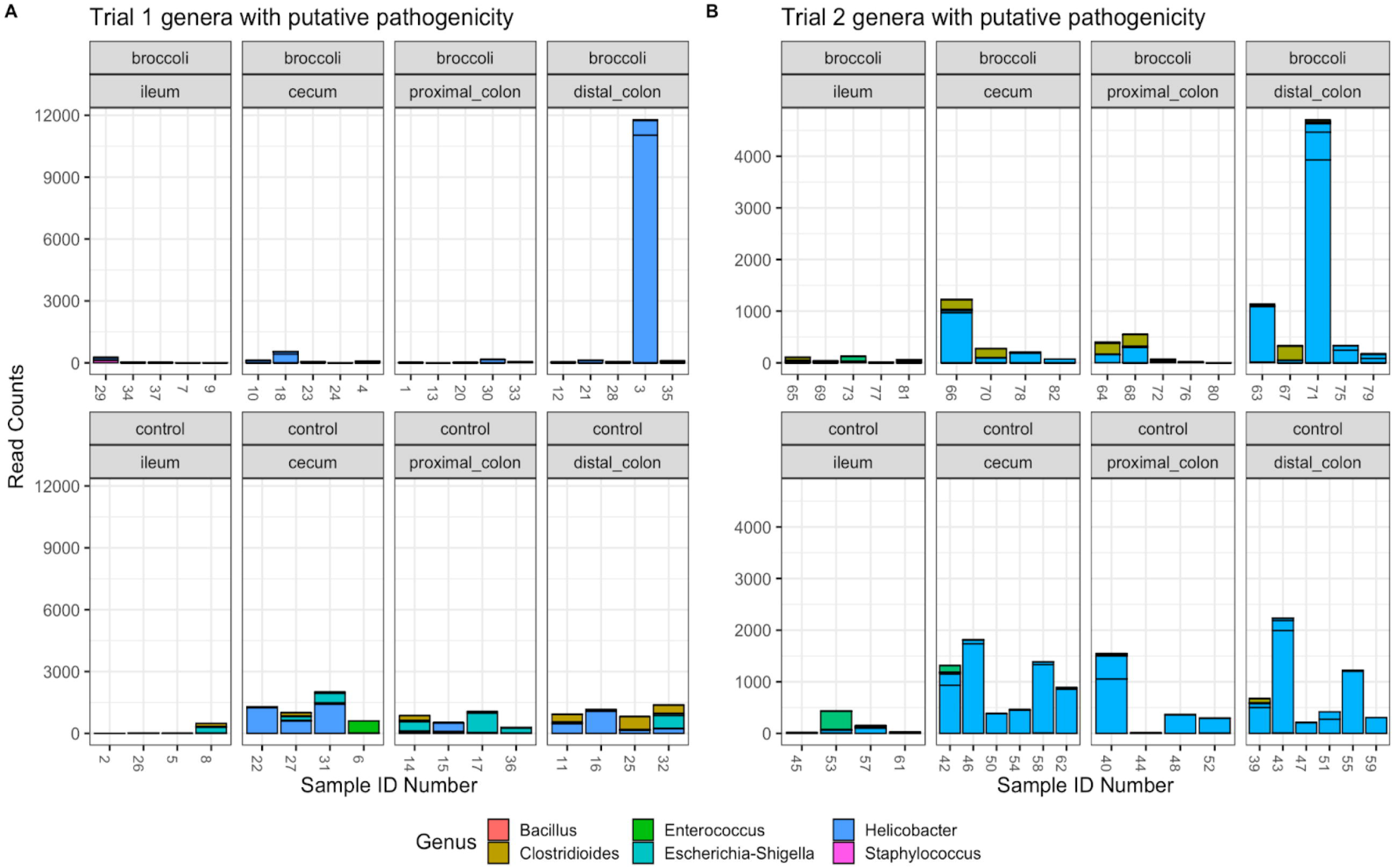
Bacterial SVs of genera which have putative pathogenicity. Trial 1 (left): 36 gut samples and 107 SVs; Trial 2 (right): 39 gut samples and 103 SVs. Graphic made using phyloseq and ggplot2 packages in R.

### Raw broccoli sprout diet increased certain putative GLR metabolizing taxa

Several bacterial taxa perform myrosinase-like enzymatic activity and metabolize GLR into SFN, and after consulting the current literature, we identified 19 taxa to analyze in detail (38, 85, 86). From our samples, we identified 562 SVs belonging to the following genera: *Bacillus, Bacteroides, Enterococcus, Lactobacillus, Lactococcus, Pseudomonas,* and *Staphylococcus* (Figure 8). Though, we did not find *Bifidobacterium, Listeria, Pediococcus, Streptomyces, Aerobacter, Citrobacter, Enterobacter, Escherichia, Salmonella, Paracolobactrum, Proteus,* or *Faecalibacterium.* There were few previously identified putative GSL-converting taxa found in the younger mice fed the broccoli sprout diet, however, there were higher read counts for *Bacteroides* and *Lactobacillus* in the younger controls as compared with older controls or broccoli-fed groups (Figure 8, ANOVA p < 0.01). Both diet groups of older mice exhibited high read counts of *Bacillus* and *Enterococcus* SVs (Figure 8).

**Figure 8.**
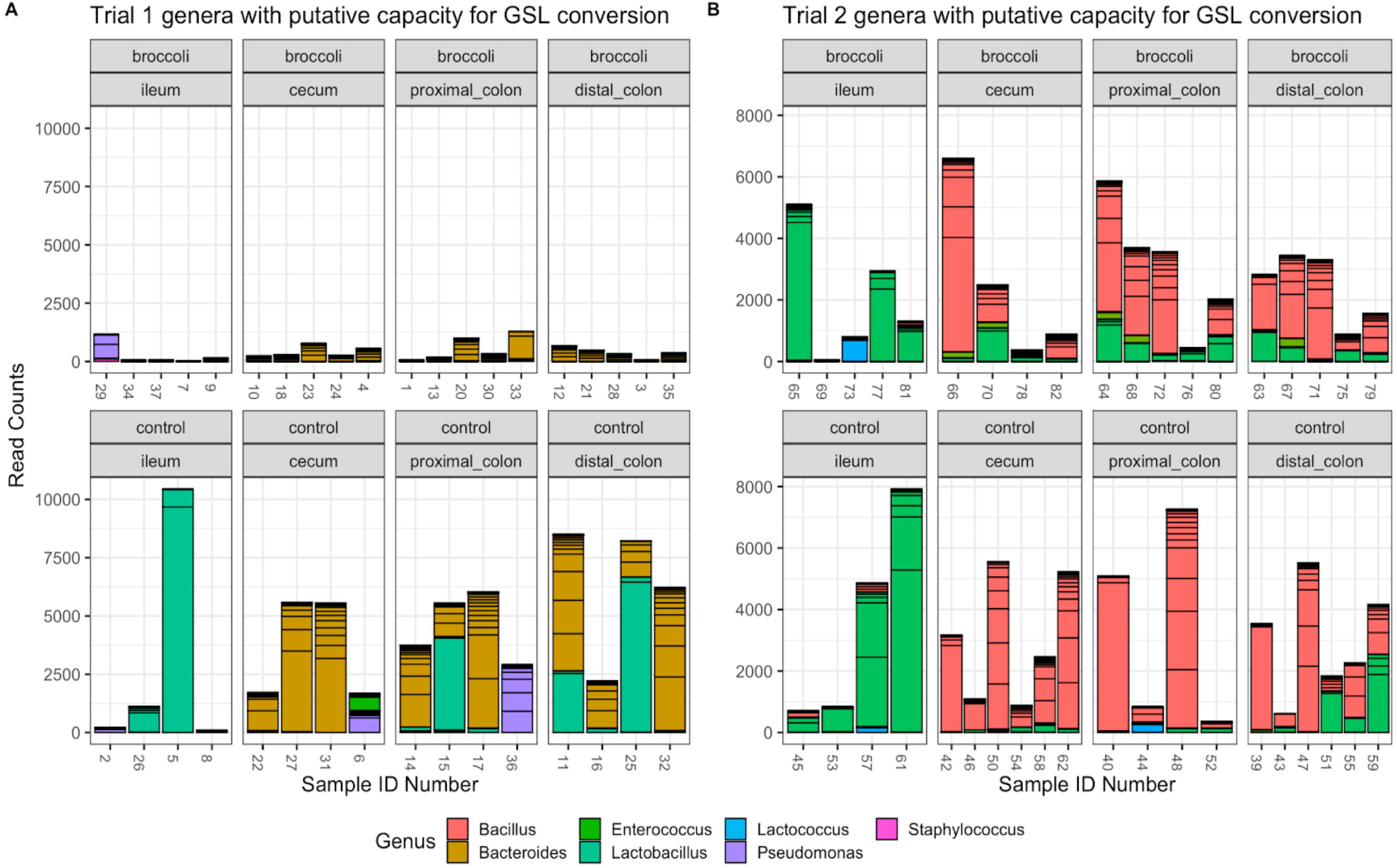
Read counts of bacterial SVs of genera which have putative capacity to convert glucoraphanin to sulforaphane (GSL). Strains of bacteria in these genera have been demonstrated to perform myrosinase-like activity in the digestive tract, as reviewed in (115). Trial 1 (left): 36 gut samples and 241 SVs; Trial 2 (right): 39 gut samples and 353 SVs. Graphic made using phyloseq and ggplot2 packages in R.

The reads from putative GSL-converting genera were identified to species-level using NCBI BLASTN (Fig. 9). The younger mice consuming sprouts contained *Bacteroides sartorii* and *B. acidifaciens* in the cecum and distal colon, and *B. caecimuris*, *B. sartorii*, and *B. acidifaciens* in the proximal colon. Most of the SVs in the ileum could not be identified to species level. In the older trial, similar GSL-converting taxa were present, with five of the identified species having high abundance (Fig. 9). The older sprout-fed mice contained many *Lactobacillus intestinalis* reads in the ileum, with fewer in the cecum, proximal and distal colon; many *B. caecimuris* and *B. acidifaciens* reads in the cecum, and proximal and distal colon; and *B. sartorii* in the proximal and distal colon.

**Figure 9.**
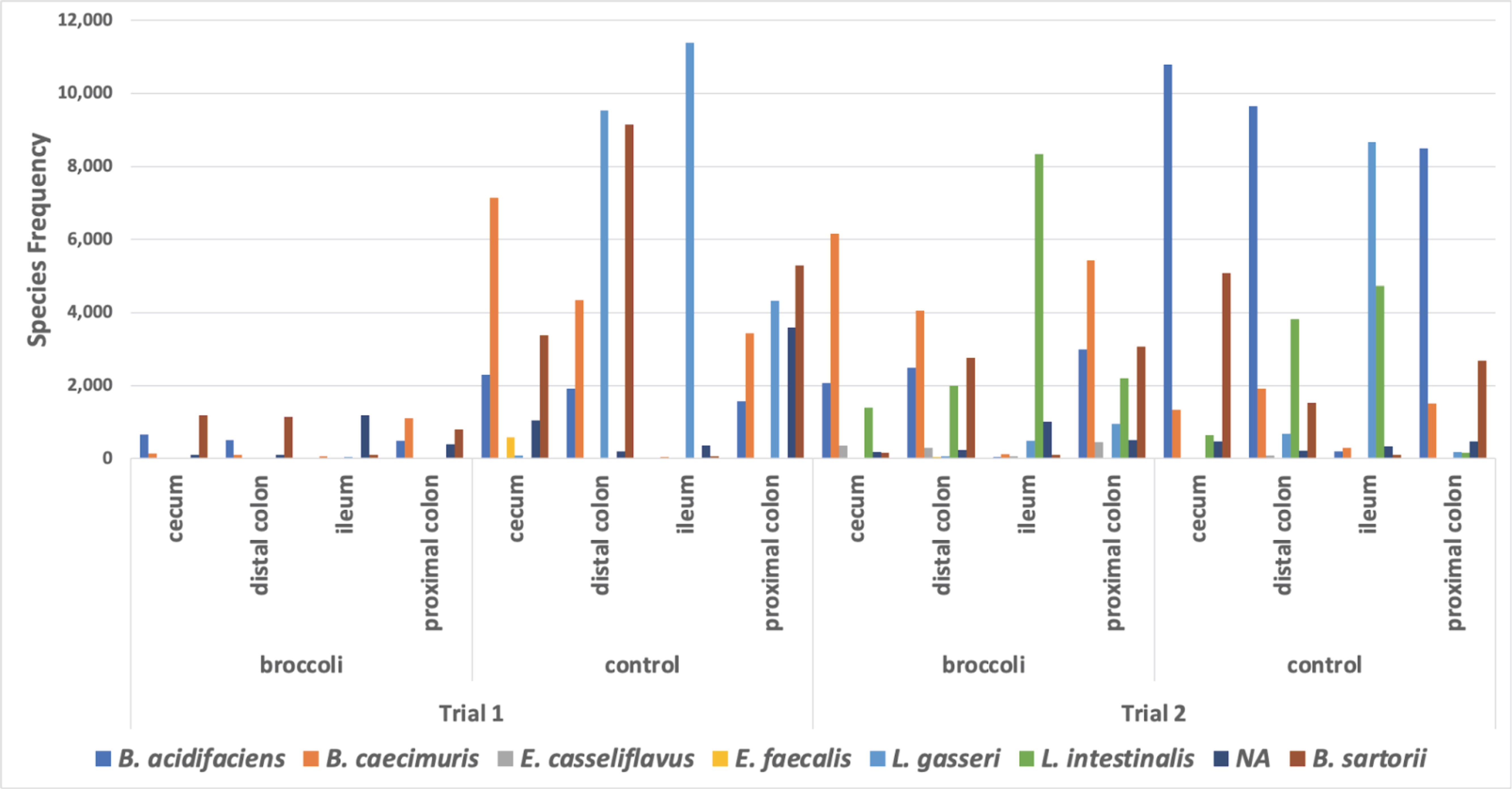
Biogeographic frequency of species from putative GSL-converting genera in the treatment groups. The Silva Database identified *Bacteroides* species *acidifaciens*, *caecimuris*, and *sartorii*, *Enterococcus cecorum*, *Staphylococcus aureus*, and *Lactobacillus intestinalis*. NCBI BLASTN identified additional *B. acidifaciens*, *B. caecimuris*, and *B. sartorii.* BLASTN uniquely identified *Enterococcus* species *casseliflavus* and *faecalis*, and *Lactobacillus gasseri*. *Bacteroides thetaiotaomicron* was linked to two (2) SVs based on genome mapping. *E. cecorum* (frequency 12), *S. aureus* (2), and *B. thetaiotaomicron* (5) were not included in the figure as their frequencies were minimal.

By contrast, the younger control group had four of the identified GSL-converting species (Fig. 9): *Lactobacillus gasseri* in the ileum, proximal and distal colon; *B. sartorii*, *B. caecimuris*, and *B. acidifaciens* in the cecum, proximal and distal colon). The older control group also had abundant *L. gasseri* in the ileum, but much less in the proximal and distal colon; *B. acidifaciens* was dominant in the cecum, proximal and distal colon; *L. intestinalis* was present in the ileum, cecum, and distal colon; and *B. sartorii* was abundant in cecum, proximal and distal colon (Fig. 9). The older control mice had a minor abundance of *E. casseliflavus*, *E. faecalis, B. thetaiotaomicron, E. cecorum*, and *S. aureus* (data not shown).

The presence of *Bacteroides thetaiotaomicron* (*B. theta*) was further investigated through qPCR amplification of its SusR4 regulon, which contains the BT2159-BT2156 operon and the regulatory gene BT2160, from samples along the digestive tract (ileum, cecum, proximal & distal colon) of mice consuming broccoli sprout and control diets. For *B. theta*, this operon has been demonstrated as necessary and sufficient for GSL metabolism and ICT production both *in vivo* and *in vitro* (*73*). These genes code for enzymes (a sugar phosphate isomerase, glycosyl hydrolase, and two oxidoreductases) that support the transformation of GLR to the bioactive SFN. We found copies of *B. theta* genes BT2156 in the ileum, cecum, and distal colon, with a small number of copies in the proximal colon (Figure S7). BT2159 had some copies in the cecum in both diet cohorts of mice, with other gene copies present in low quantities across diet and anatomical location The highest copy number of any gene, and only the significant comparison by diet, was BT2156 in the distal colon of broccoli sprout fed mice (ANOVA, difference = 92362 gene copies more than control mice, p = 0.00499, controlling for Trial).

Liou et al. found that *B. theta* sulforaphane production required BT2158 together with either BT2156 or BT2157 but was most effective when all four operon genes acted in concert (73). Thus, at each anatomical location, we characterized the distribution of the presence of the four operon genes (Figure 10A). For the young mice (Trial 1) across all anatomical sites, we found that the broccoli diet on average shifted the proportion of mice with a high operon indication (max operon mean copy count > 10,000) from 24% in controls to 64% in the treatment group (Figure 10B). For the older mice (Trial 2), the impact of the broccoli diet was not as prevalent across anatomical sites, but was indicated in the cecum and distal colon shifting the combined middle and high operon indication (max operon mean copy count > 100) in the ileum from 83% in control mice to 100%, from 66% in control mice to 80% in the cecum, and from 83% in control mice to 100% in the distal colon (Figure 10A).

**Figure 10:**
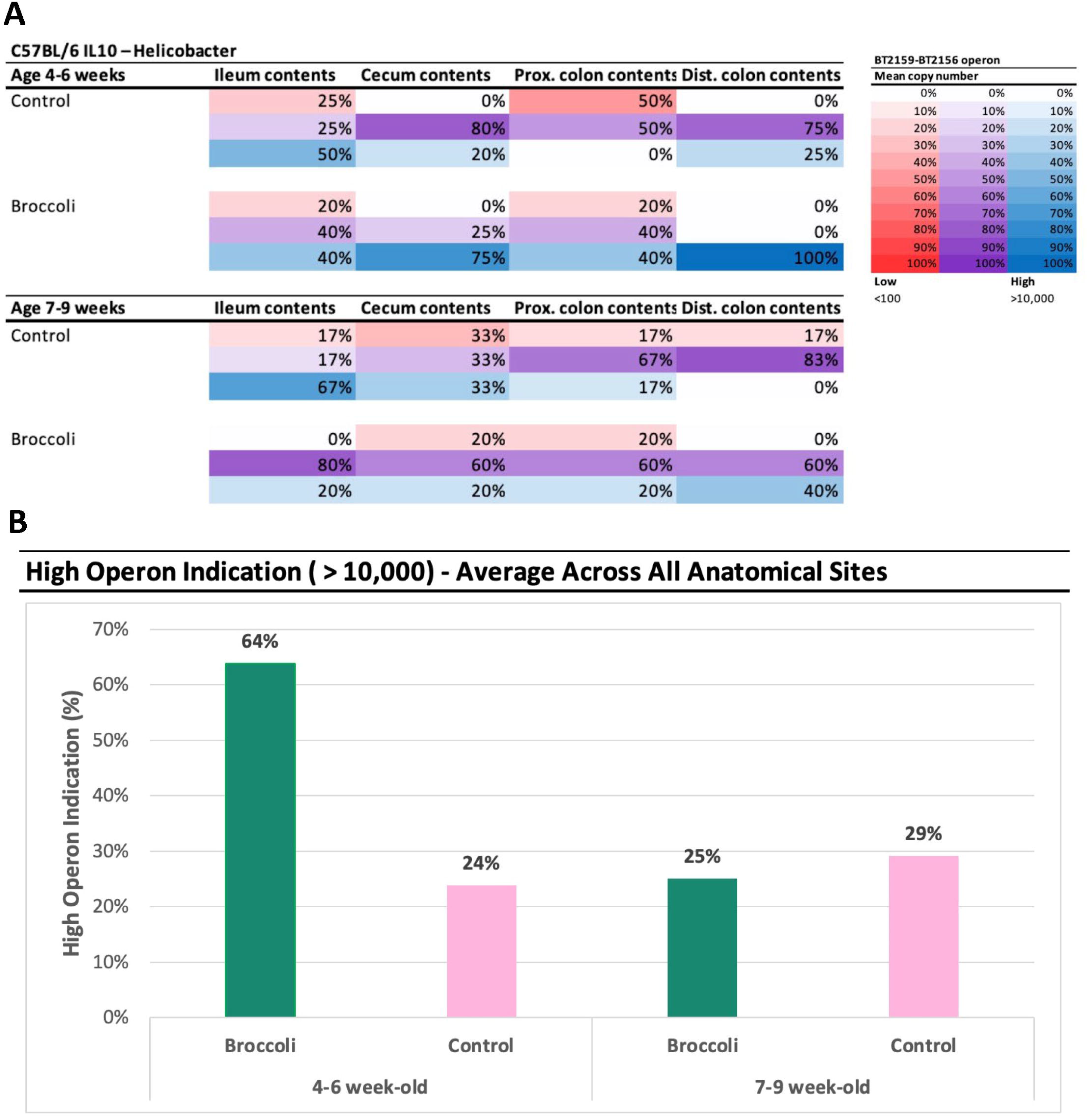
**Prevalence of the operon BT2159-BT2156 in *Bacteroides thetaiotaomicron* (VPI-5482) in 4-6 week-old (Trial 1) and 7-9 week-old (Trial 2) IL-10-KO mice fed either a control or raw broccoli sprout diet**. A. Percentages indicate the number of samples with a max operon mean copy count in each category (red low <100, purple medium 100-10,000, blue high >10,000). B. Average high indication across all anatomical sites by trial and diet.

## Discussion

### IL-10-KO is an effective model for evaluating host, diet, immune response, and microbiota

This pilot study evaluated IL-10 deficient mice as a model for using dietary broccoli sprout bioactives to reduce inflammation, modify the immune response, and support GI tract microbiota. Inflammatory diseases such as CD are associated with changes in the presence, abundance, and functionality of someone’’s microbiome (56, 57, 74), and therapies which improve the functionality of the gut microbiome have the potential to mediate other symptoms (37, 45). Specific pathogen free IL-10-KO mice acquire gut microbiota as they age (75), but they never fully develop diverse, commensal gut microbiota and this appears to be attributable to the spontaneous generation of enterocolitis at the 4 – 6 weeks of age (76). Diet is an important driver of microbiota development in early life (16), and consumption of fiber-rich foods has been associated with microbiota stabilization (77), because the wealth of microbial byproducts is demonstrated to recruit and maintain a diverse microbial community (78), reduce inflammation, improve nutrition, and stimulate gut motility – aspects which are limited in IBD (56, 79, 80). In our study, raw broccoli sprouts in the diet provided during the critical period of microbiome stabilization and enterocolitis development in these IL-10-KO mice resulted in an increase in microbial diversity, increased abundance of potentially beneficial bacteria, and a reduction of potentially pathogenic bacteria. We did not see an improvement in histological damage, which may have been precluded by the use of raw sprouts: feeding pure SFN results in high absorption in the stomach which decreases over the GI tract (81).

Dextran sodium sulfate (DSS) is an established and widely used model for studying SFN and prevention of IBD in animals (45, 52, 53), and results in a disease profile similar in progression and morphology to human Ulcerative Colitis (52, 82). DSS modifies the expression of tight junction proteins in intestinal epithelial cells, leading to a leaky epithelial barrier (83), goblet cell depletion, erosion, ulceration, and infiltration of neutrophils into the lamina propria and submucosa (84), triggering the innate immune response (85, 86). However, Bhattacharyya et al. reported DSS results in an excess of ROS species which quickly dephosphorylate Hsp27 resulting in direct IkBa dissociation, circumventing the canonical inflammatory pathway (87). As DSS instigates colitis through chemical and physical damage, without a direct interaction on the immune system, it is inadequate for investigating the specific immunohistopathology present in Crohn’s Disease. While a few studies have used IL-10-KO mice to study the effects of certain mineral additives (88, 89), isomaltodextrin supplementation (90), or high-fat diets on colitis (91, 92), the use of this model for diet studies is still nascent. We demonstrated that a medium-high concentration of raw broccoli sprouts in the diet of IL-10-KO mice induced increased bacterial richness, SFN in the plasma, and collectively reduced the symptoms of enterocolitis.

### Younger IL-10-KO mice were more responsive to diet compared to adolescent counterparts

We observed a previously unreported effect of age in early life – a three week difference – on the heightened response of younger mice to broccoli sprouts. The gut microbiota and immune systems in younger mice were more responsive to raw broccoli sprouts, and we hypothesized that this was driven by the instability of the developing gut microbiota and its amenability to selective pressures (68, 93, 94).

Mice wean at approximately 20-22 days of life (3 weeks) which is a critical period for the co-development of the immune system and gut microbiota, and is coordinated by immune factors in milk, growth of the GI tract, and improvement of epithelial barrier function (95). By 28 – 35 days of life (4 – 5 weeks), juvenile mice are beginning sexual development and still have a changing gut microbiome (96). By days 35 – 42 (5 – 6 weeks), they begin adolescence (97) during which their gut microbiome stabilizes unless it is perturbed (96). Thus, our younger mice were still in a transitional state of life and older mice were just beginning a period of life marked by stability of the immune system and gut microbiome. Our findings are consistent with previous human studies that suggest that the gut microbiota of infants and children are more plastic than that of adults (16): starting around age 3, gut microbiota fall into long-term patterns driven by consistency in diet and lifestyle (16, 77, 98, 99). Collectively, this may indicate that dietary interventions to restructure the microbiome, would be more effective in children and adolescents than in adults, which is supported by other literature (37).

Also, compared to the controls, the younger mice fed a diet containing broccoli sprouts had more bacterial genera meet the ad hoc core cutoff for prevalence and abundance. The gut microbiomes of pediatric CD patients show divergence from healthy siblings early in the disease and a more definable “disease state microbiome” than adults with CD. We found that the older broccoli-fed mice had fewer genera that met the core standard, which could suggest that the core gut microbial community diminishes with age and interventions would need to be individualized. In adults, undergoing dietary change is one of the few external factors that can “destabilize,” or subject the adult gut microbiota to change (16, 99), which may be useful in removing gut microbial communities which are not providing functional benefits and recruiting a different community instead.

### Biogeographical patterns emergent in younger mice consuming broccoli sprouts

Anatomical location within the gut, including different organs and solid-associated, liquid-associated, and mucosal-associated locations, select for different bacterial communities due to environmental conditions and anatomical features specific to each location, i.e. biogeography (100–102), even within the first few days of life (103). In IL-10-KO mice, which are raised in pathogen-free conditions and acquire some gut microbiota, there is no biogeographical signal in the gut, and the damage to the epithelium caused by inflammation or the lack of IL-10 may preclude the ability of a typical biogeographical signal to form. The addition of a conventionally raised mouse to the IL-10-KO mouse cages, and placement in a conventional mouse room, is enough to stimulate microbial transmission (104) and for IL-10-KO mice to acquire new gut microbiota even as they react negatively to them.

The third experimental factor of this study, and the one that showed to be the least significant driver of microbial community dissimilarity, was the anatomical location of sample derivation. We found that the number of bacterial taxa (SVs) in the cecum and proximal colon of the younger trial 1 mice fed broccoli sprouts was significantly elevated, as compared to the control trial 1 mice, and significant community dissimilarity between the broccoli sprout diet group and the control group in both the cecum and the distal colon. There were no significant dietary treatment effects on richness or bacterial community similarity in any anatomical location studied in the older trial 2 mice (Figure 4B). The cecum, a muscular and oblong pouch emerging from the junction of the ileum and proximal colon, houses a diverse community of fibrolytic bacteria, and specialized peristaltic movements allow for fiber in the ileocecal junction to be pulled into the cecum, as well as separate large and small fibers to allow for retention of particles to induce more microbial fermentation (105, 106). Mice have a large cecum, even in IL-10-KO mice this organ acquires a diverse microbiota (75, 107). Studies on IBD models report significant inflammation and changes to cecal microbiota in mice (45, 108, 109). It is likely that given the short duration of this experimental period, the cecum and parts of the colon that also house diverse microbiota were the only organs which were suited to recruit differential bacterial communities fast enough to be statistically significant. While humans have a small cecum relative to our size, and do suffer inflammation there in IBD patients, the effect on IBD of the microbiota there is unknown.

### Raw broccoli sprouts encouraged some putatively beneficial taxa

CD has specifically been associated with a reduction in *Faecalibacterium prausnitzii*, a putative glucosinolate metabolizer, and reintroduction of this bacteria resulted in decreased inflammation (110). However, many gut microorganisms are only useful to the host in specific metabolic conditions (111, 112), which can obfuscate our understanding of which microorganisms may be useful to try and maintain in order to preserve beneficial interactions with the host. Further, commensalism cannot be assumed for any taxa, as IL-10-KO mice infected with *Helicobacter* also exhibited higher abundance of non-commensal *Bacteroides* and *Lactobacillus* species (113).

SFN is bacteriostatic against bacterial taxa which may provide some benefit to mice, including *Bacteroides* (*114*), a genera containing known glucosinolate metabolizers (73), and non-pathogenic *Bacillus* or *Enterococcus*. Across many of our experimental groups, we found several bacterial genera, including *Bacteroides* (but not *Bacteroides thetaiotaomicron*), *Enterococcus casseliflavus*, *Lactobacillus gasseri*, and others in very low abundance, that may perform myrosinase-like activity in the digestive tract (115). *Enterococcus casseliflavus* are gram-positive, motile facultative anaerobes sourced from vegetables, with high glucosinolate degradation efficiency (116). One SV was matched to *E. casseliflavus* based on BLASTN results in the older mice, and favored the broccoli mice (1164 count) compared to the control (113 count). Although GLR was low in the broccoli diet (4 μg per gram of diet sampled), *E. casseliflavus* may have been responsible for some of the conversion to SFN.

*Lactobacillus gasseri* was the most abundant species with putative myrosinase-like enzymatic activity. A probiotic strain with implications for fighting pathogens, *L. gasseri* has hydrolyzed GLR to SFN in the cecum of rats (39, 117). Four SVs were mapped to *L. gasseri* using BLASTN. These sequences were most abundant in the control for both trials, with the highest counts occurring in the younger control mice.

Contrary to expectations, except for *Bacteroides caecimuris*, *Enterococcus casseliflavus*, and *Lactobacillus intestinalis*, the species identified from GSL converting genera tended to be found in control treatments for both age groups of mice, although it is important to note that the use of 16S rDNA precluded any assessment of microbial activity. There was a greater abundance of putative GSL converting taxa associated with the older mice (7-9 weeks-old), except in the cecum where the abundances were higher for the younger mice. The high concentration of SFN in the diet, and low concentration of the GLR precursor, may have precluded the need for metabolizing-competent bacteria, and it may be that the taxa present in younger mice were sensitive to the presence of SFN while the community present in older mice was not. In contrast, mice on the broccoli sprout diet overall demonstrated higher indications of the presence of the operon BT2159-BT2156, suggesting *B. theta* may be the primary GLR convertor in these mice, and suggesting that 16S rDNA taxonomic identification on its own may not be an accurate metric for presumptive GLR conversion in the gut. Using the presence of the operon as a metric, the use of a broccoli diet to recruit functional *B. theta* was most strongly indicated for young mice (4-6 weeks-old), supporting our *a posteriori* hypothesis.

### Raw broccoli sprouts reduced putative proinflammatory bacteria

The causal relationship between potentially pathogenic bacteria and the onset of IBD symptoms remains murky (56, 118), with postulations about the involvement of various microorganisms, including *Mycobacterium avium* subspecies *paratuberculosis*, adherent-invasive *Escherichia coli, Mycoplasma* spp., *Helicobacter pylori*, and *Chlamydia* spp. However, the role of proinflammatory bacteria in exacerbating flare-ups is clear, and proinflammatory bacteria have been identified in and found to be more abundant in IBD patients as compared to healthy children (34) or adults (56), although often newly-diagnosed children exhibit similar fecal microbial communities compared to healthy siblings (119).

SFN induces colonocytes to produce several antimicrobial compounds (120), something which is lacking in CD patients with abnormally functioning Paneth cells (121). SFN itself is bacteriostatic against several bacteria, including *Escherichia coli*, *Klebsiella pneumonia*, *Staphylococcus aureus*, *S. epidermidis*, *Enterococcus faecalis*, *Bacillus cereus,* and *Helicobacter* spp. (48, 49, 72). Interestingly, our results suggest that consumption of broccoli sprouts may prevent the accumulation of putatively harmful, pro-inflammatory bacteria. When compared to the controls, the broccoli-fed mice exhibited less prevalence and abundance of *Helicobacter*, which is a non-harmful commensal in conventionally raised and immune competent mice (122), but acts as a pro-inflammatory bacterial genus in IL-10-KO mice with little ability to moderate their own response to the presence of gut microorganisms (64). The specific species of *Helicobacter* which colonies IL-10-KO mice can also affect the host’s immune response and severity of symptoms (123).

### Whole food strategies provide multiple mechanisms of benefit and harm reduction

In many but not all people or circumstances, high-fiber diets increase microbial diversity, recruit beneficial symbiotic species in the gut, and reduce inflammation by providing antioxidants against reactive oxygen species or antimicrobials against microbial competitors (120, 124). Dietary fiber has the potential to reduce the risk of Crohn’s flare-ups by up to 40% (125), and high-fiber foods can stimulate beneficial microbial byproducts in the gut (124). However, during a flare-up, high-fiber foods can worsen symptoms, and certain fiber types can exacerbate symptoms in IBD patients (126). Opting for foods rich in soluble fiber, which can help slow down digestion and ease diarrhea, is recommended for individuals with Crohn’s disease. On the other hand, foods containing insoluble fiber can increase water content in the gut, leading to rapid digestion, watery diarrhea, stomach cramps, or gas, and in severe cases, may cause blockages. Using purified GLR or SFN have been evaluated as a means of inducing health benefits while avoiding fiber in IBD patients, with varying effectiveness, e.g., (40, 51).

Feeding SFN directly to mice, thus eliminating fiber risk, does not appear to change cecal butyrate (127), and while butyrate treatments to the gut are often successful in patients with ulcerative colitis it is unclear whether butyrate is helpful for patients with CD as cellular damage may prevent colonocytes from utilizing it (128). We balanced the 10% raw broccoli sprout/90% control diet to the 100% control diet by total fiber, and the control diet contains oats which are a prime source for microbially-produced butyrate (129). While mature broccoli is a reliable source of fiber and stimulates butyrate production by gut bacteria, there is no literature to suggest that broccoli sprouts would stimulate more microbial fermentation and byproducts than the equivalent grams of fiber from oats. Further, we found no putative butyrate-producing genera in our sprout-fed mice: no *Bifidobacterium*, *Clostridium*, or *Butyrococcus*, and *Bacteroides* were only high in the younger control mice. Similarly, *Bifidobacterium*, *Clostridium*, and *Bacteroides* can dissociate bilt salts in the colon, which otherwise exacerbate symptoms in IBD patients, and we found *Lactobacillus intestinalis,* also capable of this (130), in our sprout-fed mice. Finally, mucin degrading capacity of gut bacteria is reduced in CD and UC patients (131), and alterations to the gut microbiota could have action here.

Although some studies have indicated positive outcomes with anti-inflammatory dietary interventions in children and adolescents with Crohn’s disease, their effectiveness may vary among individuals. Factors such as adherence to the prescribed diet, individual tolerance to specific foods, and the involvement of healthcare professionals in closely monitoring progress all play critical roles. Diet can also trigger symptoms of IBD, however; It is important to note that there is no singular food or food group that can be universally linked to every case of Crohn’s disease, as different individuals may experience gut irritation and inflammation from different foods. Children and adolescents with Crohn’s disease may exhibit distinct manifestations compared to adults, often presenting with more extensive disease involvement, including perianal disease and structural complications (34–36). Consequently, personalized recommendations are often needed, and the unique nutritional requirements of pediatric and adolescent populations must be addressed to ensure adequate nutrient intake for growth, development, and overall health (37).

### Limitations and Future Directions

Due to the short duration of the experiment, we were unable to discern if the increase in diversity persists into adulthood, and if this would preclude the development of more intense symptoms and damage to the intestinal epithelium. We lacked sufficient power to determine sex specific differences in response to diet (90, 132), immune function (133), and microbial acquisition (134). We also lacked data to determine microbial activity, beneficial or detrimental.

Future research will be required to examine the effect of cooking preparation on the concentrations of precursors and bioactives which are contained in the diet and available in the gut, as well as microbially sourced bioactives, and will require metabolomics to fully explore this. The myrosinase enzyme present in broccoli and sprouts can metabolize the GLR precursor to bioactive SFN when the plant tissues are cut or chewed, but most of the precursor is converted to biologically inactive SFN nitrile by the epithiospecifier protein, also present in broccoli (14). SFN was high in our raw sprout diet, which was an effect of the preparation process stimulating release of myrosinase. However, SFN is an unstable molecule and will not persist in processed raw sprouts unless kept frozen. These enzymes can be inactivated by cooking, preserving the stable precursor in the diet and allowing gut microbiota to perform the conversion in the intestines (38).

Future research will be needed into the feasibility and adoptability of this intervention. The difficulty in managing CD symptoms is costly to patients (135), and healthcare systems (136, 137). Further, many IBD patients may be told to avoid fiber for fear of exacerbating symptoms or aggravating tender intestines (24, 54). Given the importance of diet in supporting host health and microbiota, we underscore the need for understanding dietary preferences, and for incorporating nutritional intervention as part of a holistic treatment for gastrointestinal inflammation.

## Materials and Methods

### Diet

Multiple lots of Jonathan’s Sprouts™ (Rochester, MA, U.S.) broccoli sprouts were purchased from a nearby grocery store (Bangor, ME, U.S.) and stored in a –80L freezer until freeze-drying at the University of Maine Pilot Plant (Orono, ME, U.S.) to lower moisture content and reduce enzymatic activity. The control diet was 100% irradiated 5LOD diet irradiated 5LOD rodent diet powder (ScottPharma Inc., Marlborough, MA, #50795) mixed with ultra-pure water. The freeze-dried sprouts were crushed by mortar and pestle into a fine powder and mixed with the 5LOD powder and water to a concentration of 10% by weight to form the treatment diet, as that concentration reliably produces consistent anti-inflammatory results (46). Diet pellets were formed using a silicone mold to ensure consistent sizing, dried at room temperature for up to 48 hours in a chemical safety hood to facilitate moisture evaporation, and after drying were stored in Ziploc bags in a –10L freezer until future use. The sprouts are washed prior to packaging, and freezing, freeze-drying, and drying reduces microbial biomass, growth, and activity, and may kill some bacteria through desiccation. However, we did not verify that diets were sterile after formulating them, as the goal of the experiment was to introduce microbiota to mice.

The fiber content of the control diet and broccoli sprout-supplemented diet was ∼5%. No GLR or SFN was found in the control and the sprout diet contained an average 4 μg of GLR and 85 μg of SFN (induced when raw diets are crushed and myrosinase is released from broccoli tissue vesicles) per gram of dried diet sampled (Figure S8), using LC/MS (38).

### IL-10 Mouse Model

All experimental protocols were approved by the Institutional Animal Care and Use Committee of the University of Vermont (PROTO202000040), in Burlington, Vermont, U.S. and the Institutional Biosafety Committee of the University of Maine (protocol SI-092220). Two replicate trials were conducted using male and female IL-10-KO mice (*Mus musculus*) on a C57BL/6 background (B6.129P2-Il10tm1Cgn/J, strain 2251; Jackson Laboratories, Bar Harbor, Maine, U.S.). IL-10-KO mice are a well-demonstrated model for Crohn’s research and the C57BL/6 genetic line shows some genetic-based resistance to developing symptoms (61), which allowed us to focus on environmental triggers of disease.

Trial 1 included 9 mice (n = 4, control; n = 5 sprout) starting at 4 weeks of age and post-weaning. Trial 2 included 11 mice (n = 6, control; n =5 sprout) starting at 7 weeks of age. The experimental design was structured as a prevention paradigm (Figure 1). A colony of homozygous IL-10-KO mice was maintained in a barrier facility which is specific pathogen free, and especially *Helicobacter (H.) hepaticus* free. For the duration of the trial, they were fed either the 5LOD control diet or the treatment diet consisting of 10% (w/w) raw broccoli sprouts and 90% control diet, which was balanced for micronutrients and fiber (5% of diet). Mice had access to tap water *ad libitum*.

Mice were acclimated to diets for 7 days prior to the induction of colitis, which was achieved by moving mice to a conventional mouse room and adding an *H. hepaticus* positive mouse in the cage for the remainder of the experiment. The *H. hepaticus* positive mouse transfers commensal bacteria, as well as species which may cause infection in immunocompromised mice, via feces and grooming, and bacteria induce colitis over the course of 10 days (61, 67). While *H. hepaticus* is widespread in research mouse colonies and causes disease in immunocompromised mice, it does not cause symptoms in healthy, conventional mice (122). The conventional mice were fed an irradiated RMH 3000 diet (Prolab® IsoPro®) prior to being co-housed with IL-10-KO mice, at which point they consumed the control or 10% broccoli sprout diets based on the cage treatment. Day 0 of each trial was set as the day the IL-10 mice were exposed to the *H. hepaticus* positive mouse, such that experimental Day 10 is when symptoms appear (Figure 1). Animals were euthanized by carbon dioxide and cervical dislocation on Day 16.

### Assessment of Disease Activity Index

Disease severity was assessed every other day beginning on Day 0 when mice were co-housed with the *H. hepaticus* positive mouse, using a DAI. This included a score that combined fecal blood presence, severity, and stool consistency (138). The animals’ weights were normalized to their baseline weight on Day 0 of the trial, as at this age, mice are still in a growth phase and likely to gain weight even when under duress (139); and this was the rationale for excluding weight loss as part of the DAI criteria. The presence of fecal blood was assessed using Hemoccult Single Slide testing slides (Beckman Coulter, Brea, CA, U.S.).

### Assessment of Lipocalin and Proinflammatory Cytokines

Plasma samples collected immediately after euthanasia, as well as fecal samples at various time points, were frozen for evaluation of fecal lipocalin (LCN2); a neutrophil protein that binds bacterial siderophores and serves as a biomarker for intestinal inflammation (71). Frozen fecal samples were weighed up to 20 mg and reconstituted in phosphate buffered solution (PBS) with 0.2 mL of 0.1% Tween 20, thawed, and vortexed to create a homogenous suspension. The samples were centrifuged for 10 minutes at 12,000 rpm at 4 °C. Clear supernatant was collected and stored at –20 °C until analysis. LCN2 concentration was measured by a mouse Lipocalin-2/NGAL DuoSet ELISA kit (R & D Biosystems, USA) following the manufacturer’s instructions. The readings at wavelengths of 540 nm and 450 nm were measured by a Thermo Scientific Varioskan LUX Multimode Microplate Reader. The readings at 540 nm were subtracted from the readings at 450 nm to correct for the discoloration of the solution in serum.

We analyzed several pro-inflammatory cytokines in the mouse plasma samples which were collected at the end of the study: CCL4 /macrophage inflammatory protein 1β (MIP1β), which regulates the migration and activation of immune cells during inflammation (140); and IL-1β, IL-6, and tumor necrosis factor alpha (TNF-α), which play several roles regulating immune responses (141–143). The concentrations of mouse CCL4/MIP-1β, IL-1β/IL1F2, IL6, and TNF-a were analyzed using the Simple Plex Ella Automated Immunoassay System (Ella) from R&D Biosystems, USA. Mouse serum samples were diluted 10-fold using the kit-specific reagent (SPCKA-MP-007374, Protein Simple, Bio-Techne), and the concentrations were measured following the manufacturer’s instructions on the Ella system. Mean values were calculated for each cytokine and analyzed using ANOVA.

### Quantification of Plasma SFN

Plasma collected after euthanasia was also used for SFN measurement using LC/MS (38). To extract SFN, 50 µl of plasma homogenate was mixed with 3-5 fold of acetonitrile and centrifuged at 14,000 rpm, 4 °C for 15 min, and the supernatant stored at –20°C. An AB SCIEX QTRAP 4500 with a TurboV electrospray ionization source mass spectrometer (Applied Biosystems, Carlsbad, CA) coupled to an Agilent 1200 Series HPLC system (Agilent Technologies, Santa Clara, CA) was used for the quantification. HPLC separation was performed on a Waters XBridge® C18 3.5 μm 5 cm x 2.1 mm column (Waters Corporation, Milford, MA). Mobile phase A (water containing 0.1% formic acid) was first kept at 90% for 0.5 min, and then decreased to 5% over 1 min and maintained at 5% for 2 min, and then returned to 10% mobile phase B (ACN containing 0.1% formic acid) and maintained for 3 min. The flow rate was 400 L/min. Positive ion MS/MS was conducted to detect SFN with the following conditions: source temperature, 400 °C; curtain gas (CUR), 30 psi; ionspray voltage (IS), 4500 V; desolvation gas temperature (TEM), 500 °C; ion source gas 1 (GS1), 60 psi; ion source gas 2 (GS2), 40 psi; collision gas (CAD), high; entrance potential (EP), 4 eV; collision energy (CE), 15 eV. The MS/MS transition of 178 > 114 was used to detect SFN, and a dwell time of 50 ms was used for the transition. Data acquisition and quantitation were performed using Analyst software (Applied Biosystems, Carlsbad, CA).

### Histological Analysis of Tissues

After euthanasia, tissue segments (1 cm, collected in duplicate) from the ileum, proximal colon and distal colon were collected and fixed in 4% paraformaldehyde overnight for histological evaluation. Tissues were immediately rinsed with phosphate buffer solution (PBS), placed in 2% paraformaldehyde/0.2% picric acid as a preservative, and stored at 4L until transport to the University of Maine Electron Microscopy Laboratory (Orono, Maine) for processing. All processing protocols took place in a biosafety cabinet using aseptic techniques to reduce contamination. A pipette was used to gently remove the previous PBS that the tissue was submerged in without disturbing tissue while leaving a little bit of PBS behind to prevent the tissue drying out. The sample tubes were refilled with fresh PBS, and the wash step was repeated four times throughout the day every three hours with samples stored at 4L between washes.

After the final wash step, the tissue samples were transferred into embedding baskets. The samples were dehydrated by immersing the samples in a graded series of ethanol at 4L (50%, 70%, 80%, 95%, 100% ethanol), cleared in xylene, and then infiltrated and finally embedded in Paraplast X-tra. During embedding, care was taken to orient the intestine samples to give a cross section during sectioning. Slides were sectioned at 5μm and stained in Hematoxylin and Eosin (144). Images were taken using an Olympus BX41 microscope with a Zeiss ERc 5s digital camera.

The tissues were scored with 7 criteria (Figure S1) used to assess inflammation in the ileum, and the proximal and distal colon tissues, performed according to parameters provided by the Mawe Lab at the University of Vermont (145). Epithelial damage and architectural changes were scored from 0 (no damage) to 2 (extensive damage). Similarly, infiltration of mononuclear cells in the lamina propria was scored on a scale of 0 (no infiltration) to 2 (extensive infiltration). Infiltration of polymorphonuclear cells in both the lamina propria and the epithelium were scored on a scale of 0-2, with 0 indicating no infiltration, 1 equating to sighting of ≥1 cell in each viewing field, and 2 equating to >3 cells in each viewing field. Abscesses, ulcers, erosion, and branched crypts were scored together with 0 being absence of these damage indicators and 1 being presence. Finally, presence of granulomas was scored with 0 indicating no granulomas and 1 equating to ≥1 granulomas. Scores by treatment, anatomical location, and trial were compared with analysis of variance.

### DNA extraction and 16S rRNA bacterial sequencing library preparation

On Day 16, following euthanasia, intestinal tissue segments (2 cm in length) were collected from the ileum, cecum, proximal colon, and distal colon; placed in RNAlater preservative (Invitrogen, Waltham, MA, U.S.); and transported overnight on ice to the University of Maine for DNA extraction. All DNA extraction processing steps took place in a biosafety cabinet using aseptic techniques to reduce contamination. All tissues containing their resident gut microbiota were gently homogenized with vortexing, then treated with propidium monoazide (PMA; BioTium) following kit protocols at a final concentration of 25 μmol. PMA covalently binds to relic/free DNA and DNA inside compromised/dead cell membranes, and prevents amplification in downstream protocols to preclude dead DNA from the sequence data.

Following PMA treatment, bulk DNA was extracted from tissue-associated bacterial communities (n = 80 samples), or no-template (water) control samples (n = 4, one for each extraction batch) using commercially available kits optimized for fecal-based microbial communities (Zymo fecal/soil kit), and some aliquots archived. DNA extract was roughly quantified and purity-checked with a Nanodrop spectrophotometer. Samples underwent DNA amplicon sequencing of the 16S rRNA gene V3-V4 region, using primers 341F (146) and 806R (147) and protocols consistent with The Earth Microbiome Project (148), and sequenced on an Illumina MiSeq platform using the 2 x 300-nt V3 kit (Molecular Research Labs, Clearwater, TX). Nine samples failed the first sequencing run, and were extracted again for repeated sequencing.

Raw sequence data (fastq files and metadata) from both runs are publicly available from the NCBI Sequence Read Archive (SRA) under BioProject Accession PRJNA909836.

### 16S rRNA bacterial community sequencing analysis

Amplicon sequence data was processed using previously curated workflows in the Ishaq Lab (Supplemental Material) which used the DADA2 pipeline ver. 1.22 (149) in the R software environment ver. 4.1.1 (150). The dataset started with 38,287,212 million raw reads from 80 samples and 4 negative controls. All samples were sequenced in an initial batch together, but 5 samples failed to meet all quality control standards and were resequenced. The two sequencing batches were put through quality control steps separately in DADA2 and combined before rarefaction. Trimming parameters were designated based on visual assessment of the aggregated quality scores at each base from all samples (plotQualityProfile in DADA2): the first and last 10 bases were trimmed, and sequences were discarded if they had ambiguous bases, more than two errors, or matching the PhiX version 3 positive control (Illumina; FC-110-3001). After filtering, 43,382,928 paired non-unique reads and 289 samples remained.

The DADA algorithm was used to estimate the error rates for the sequencing run, dereplicate the reads, pick SVs which represent ‘microbial individuals’, and remove chimeric artifacts from the sequence table. Taxonomy was assigned using the Silva taxonomic training data version 138.1 (151) and reads matching chloroplasts and mitochondria taxa were removed using the dplyr package (152). No-template control samples were used to remove contaminating sequences from the samples by extraction batch (153). The sequence table, taxonomy, and metadata were combined using the phyloseq package (154) to facilitate statistical analysis and visualization, representing 80 samples and 22,805 taxa. Due to the variability in sequences per sample which passed quality assurance parameters (range 12,060 – 157,040 sequences/sample), the data were rarefied (155, 156) to 12,060 sequences/sample.

Normality was checked using a Shapiro-Wilks test on alpha diversity metrics generated from rarefied data; observed richness (W = 0.86662, p-value = 1.186e-06); evenness (W = 0.89141, p-value = 9.775e-06); and Shannon diversity were not normally distributed (W = 0.95012, p-value = 0.005061). Linear models compared alpha diversity metrics (lme4 package (157)), in which anatomical location, diet treatment, and trial (either 1 or 2) were used as factors. Jaccard unweighted similarity was used to calculate sample similarity based on community membership (species presence/absence), and non-parametric multidimensional scaling (Trial 1 run 20 stress = 0.1940131; Trial 2 run 20 stress = 0.1748438) and tested with permutational analysis of variance (permANOVA) by using the vegan package (158). Random forest feature prediction with permutation was used to identify differentially abundant SVs based on factorial conditions (159). Plots were made using the ggplot2 (160), ggpubr (161), and phyloseq packages. Source Tracker algorithms which had been modified for the R platform (162, 163) were used to identify source: sink effects based on anatomical location. This was used to determine if the cecum could be the source for population sinks in the colon, as a proxy for the model’s applicability to the human gut anatomical features and microbial communities (Figure S6).

### Quantitative PCR performed on genes which metabolize glucoraphanin

The ability of gut bacteria to perform glucoraphanin conversion to sulforaphane was assessed using gene sequences and primers associated with the established pathway in *Bacteroides thetaiotaomicron* VPI-5482 (73) (see Table S1). Gene sequences of interest from *B. thetaiotaomicron* VPI-5482 were evaluated through the ApE-A plasmid Editor v3.1.3 to confirm optimal melting temperature and GC content of primers, and assessment of primers was performed using NCBI primer blast (https://www.ncbi.nlm.nih.gov/tools/primer-blast/) and IDT’s OligoAnalyzer (www.idtdna.com). Quantitative polymerase chain reaction (qPCR) was completed on extracted DNA to determine copy numbers of glucoraphanin-metabolizing genes previously established in *B. thetaiotaomicron* using a Applied Biosystems QuantStudio 6 Flex Real-Time PCR system (Applied Biosystems, Foster City, CA, USA), primer sets BT2156-BT2160, and Luna® Universal qPCR Master Mix were employed. After diluting primers to 10 µM working concentration, the primer concentration in each well was 0.25 µM. PCR conditions consisted of one cycle of 50°C for 2 min, one cycle of 95°C for 1 min, and forty cycles of 95°C for 15 sec, 60°C for 30 sec and 72°C at either 20, 25 or 30 sec based on the primer set used (Table S1). Standard curves were generated for each gene by serially diluting geneblocks (IDT) six times. All standards and samples were tested in triplicate, with standards and negative controls present on both plates. Using the standard curve produced in the Quantstudio analysis software, sample gene copy numbers were quantified. R software was used to run an analysis of variance, with corrections applied for multiple comparisons through Tukey’s HSD.

## Acknowledgements

All authors have read and approved the final manuscript. This project was supported by the USDA National Institute of Food and Agriculture through the Maine Agricultural & Forest Experiment Station: Hatch Project Numbers ME022102 and ME022329 (Ishaq) and ME022303 (Li); the USDA-NIFA-AFRI Foundational Program [Li and Chen; USDA/NIFA 2018-67017-27520/2018-67017-36797]; and the National Institute of Health [Li and Ishaq; NIH/NIDDK 1R15DK133826-01] which supported Marissa Kinney, Timothy Hunt, and Benjamin Hunt. Johanna Holman was supported by ME0-22303 (Li), and Lola Holcomb was supported by US National Science Foundation One Health and the Environment (OG&E): Convergence of Social and Biological Sciences NRT program grant DGE-1922560, and the UMaine Graduate School of Biomedical Science and Engineering.

## Author Contributions

Conceptualization, S.L.I., Y.L., T.Z., G.M., P.M., G.C.; Methodology, S.L.I., Y.L., T.Z., G.M., P.M., J.H., L.H., M.H., B.L., EP; Software, S.L.I.; Formal Analysis, J.H., L.H., S.L.I.; M.H., A.S., T.H., B.H., M.K.; Investigation, M.H., B.L.; J.H., L.H., L.C.; A.S., J.P.; Resources, G.M., S.L.I., Y.L.; Data Curation, S.L.I., J.H., L.H.; Writing – Original Draft, J.H., L.H., S.L.I., Y.L.; Writing – Review and Editing; S.L.I., Y.L., T.Z., G.M., P.M., J.H., L.H., M.H., B.L., G.C.,T.H., B.H., E.P., M.K., J.P.; Visualization, J.H., L.H., S.L.I., A.S.,T.H., B.H., M.K.; Supervision, S.L.I., Y.L., T.Z.; G.M.; Project Administration, S.L.I., Y.L., G.M.; Funding Acquisition, S.L.I., Y.L., G.M. G.C.

## Supplemental Material Legends

**Supplemental Figures and Table.** Additional data visualizations are included to demonstrate minor points or null results.

**IL10 mouse R code.** The R code that was used to clean, visualize, and statistically analyze the bacterial community sequencing data presented.

