## Supplemental Figure and Table for "Early life exposure to broccoli sprouts confers stronger protection against enterocolitis development in an immunological mouse model of inflammatory bowel disease"

**Supplemental Figures and Table.** Additional data visualizations are included to demonstrate minor points or null results.

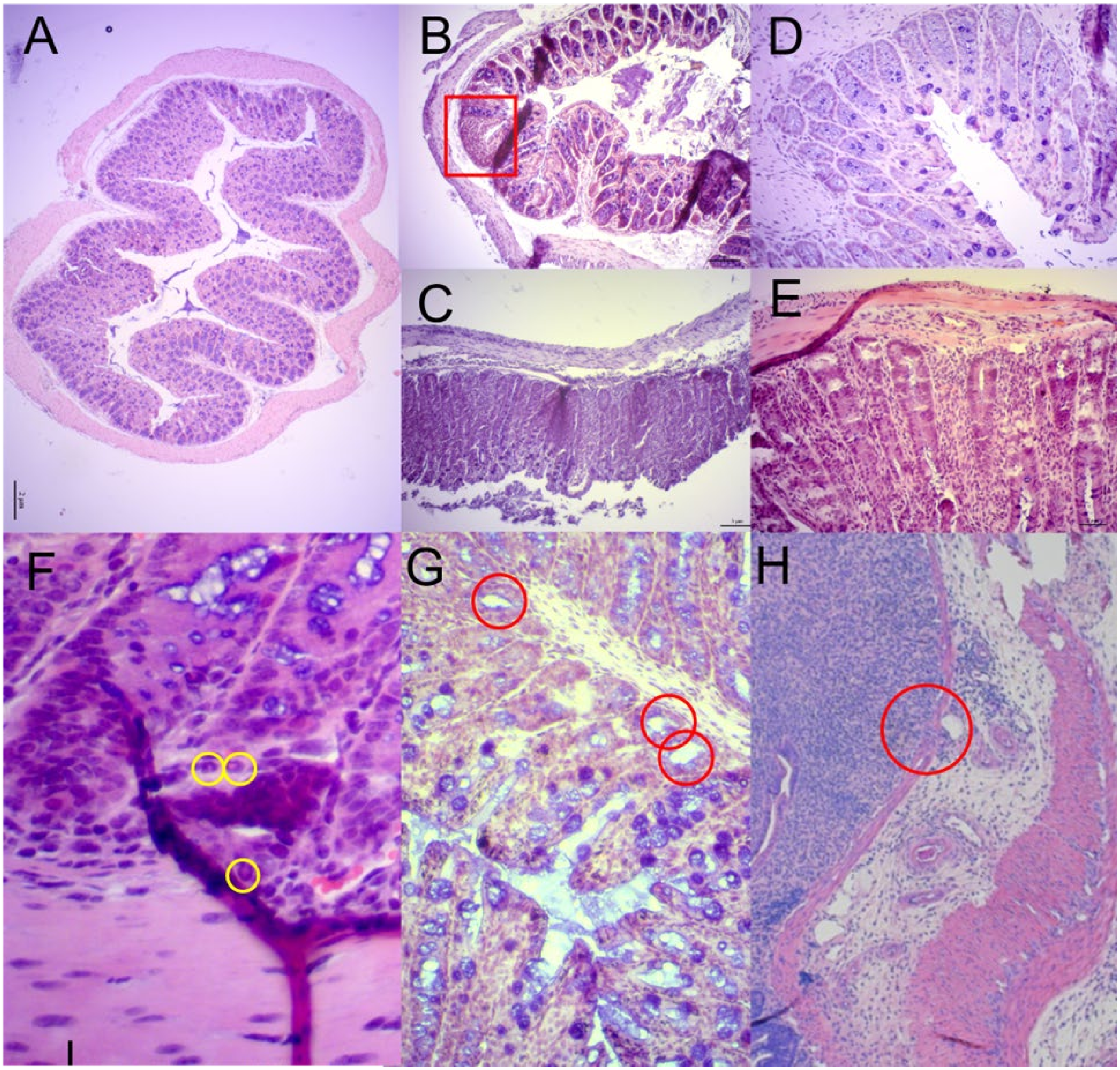


**Figure S1. Histology scoring methodology for mouse colon tissues.** A. Healthy colon. B. Focal Epithelial Damage. C. Extensive Epithelial Damage. D. Moderate mononuclear cell infiltration. E. Extensive mononuclear cell infiltration, also demonstrates architectural changes. F. Presence of multiple polymorphonuclear cells. G. Crypt abscess. H. Ulceration. Detailed protocols available in Spohn et al. 2026 Gastroenterology 151:933–944.e3.

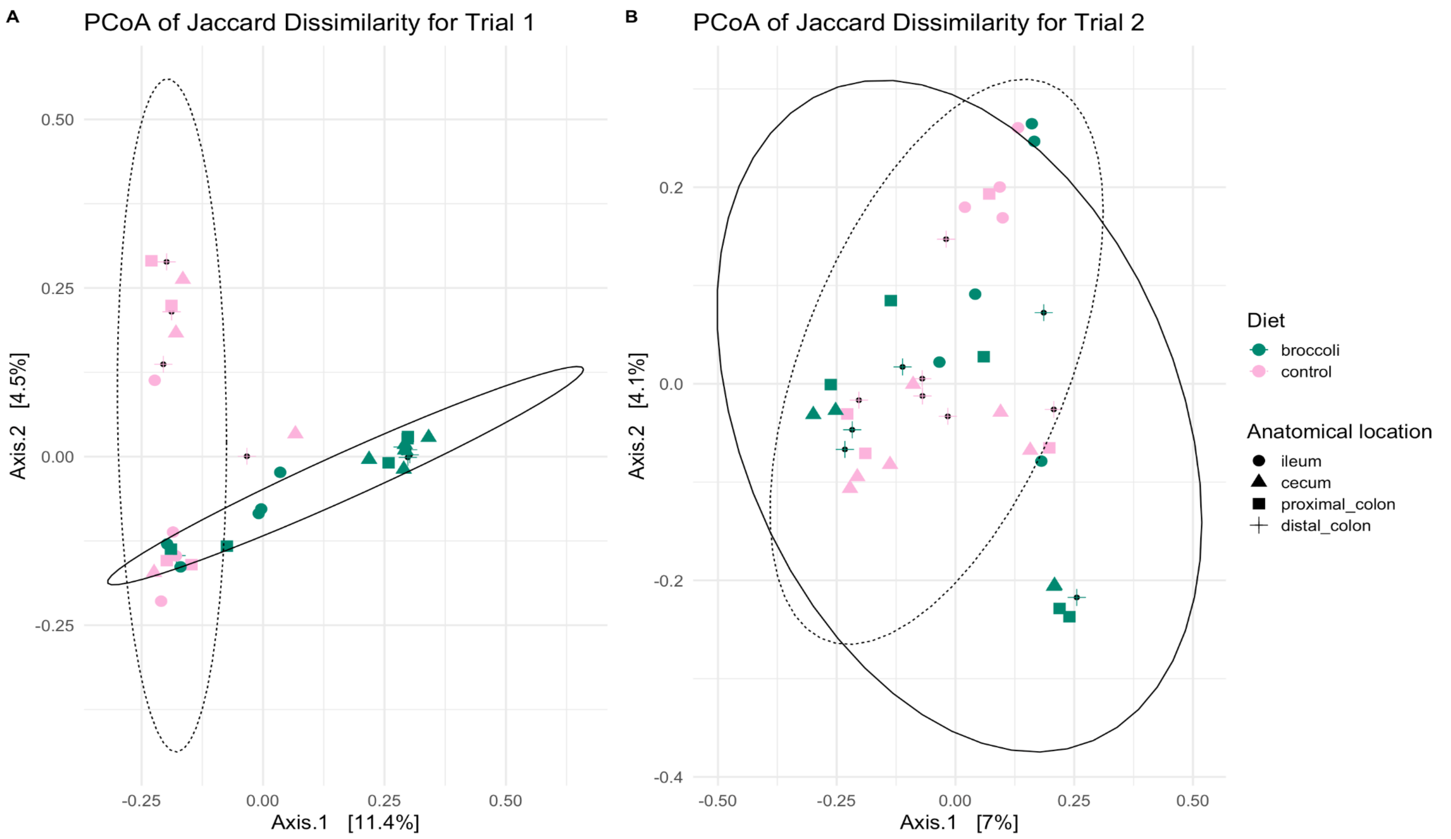


**Figure S2. Principal coordinates analysis of bacterial community similarity within the gastrointestinal tracts of 4- or 7-week-old IL-10-ko mice fed control diets or broccoli sprout diets.** Calculations using unweighted Jaccard dissimilarity show differences in the taxonomic structure of trial 1 mice in Panel A, and trial 2 mice in Panel B. Graphic made using phyloseq, and vegan packages in R.

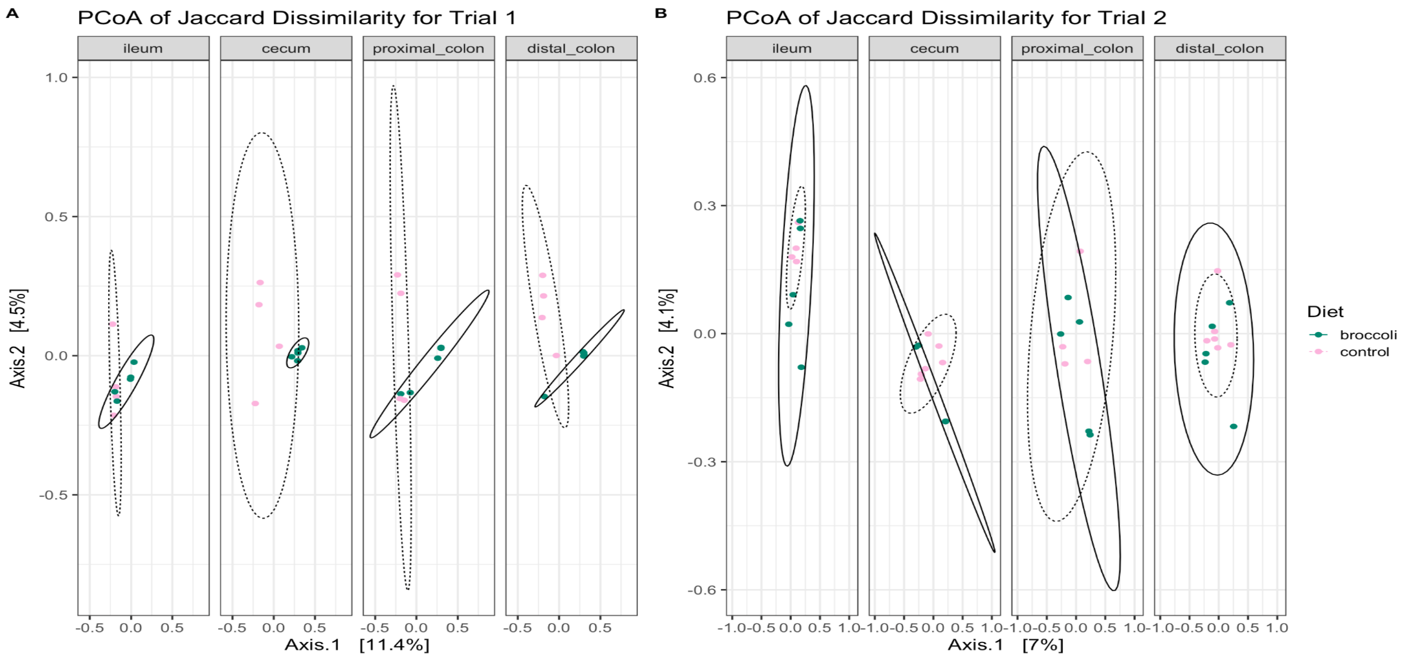


**Figure S3. Bacterial community dissimilarity within a specific anatomical location along the gastrointestinal tract of IL-10-KO mice fed control diets or broccoli sprout diets beginning at 4 or 7 weeks of age.** Calculations using unweighted Jaccard dissimilarity show differences in the taxonomic structure of (A) trial 1 mice sampled at age 6 weeks, and (B) trial 2 mice sampled at age 9 weeks, after each had been consuming diets for 3 weeks. Graphic made using phyloseq, and vegan packages in R.

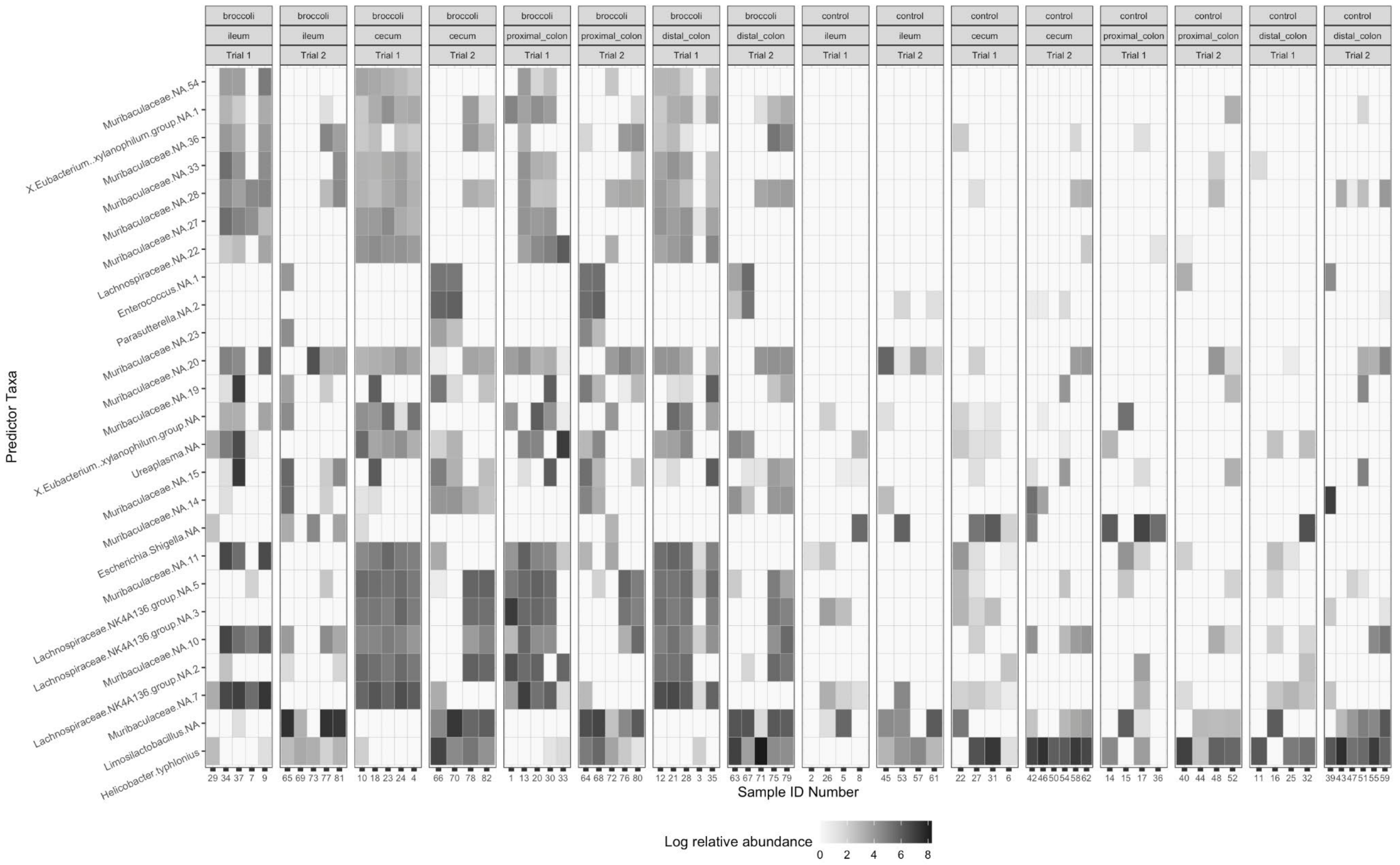


**Figure S4. Abundance of bacterial SVs identified as differential from all experimental factors.** Top 25 SVs were identified through permutational random forest analysis. Random forest classification (number of trees: 500, number of variables tried at each split: 98, number of permutation replicates: 100) model accuracy: broccoli percent correct: 61.5%, control percent correct: 100%, overall percent correct: 80%. 96 significant SVs identified. Graphic made using phyloseq, vegan, plyr, dplyr, randomForest, rfPermute, and ggplot2 packages in R.

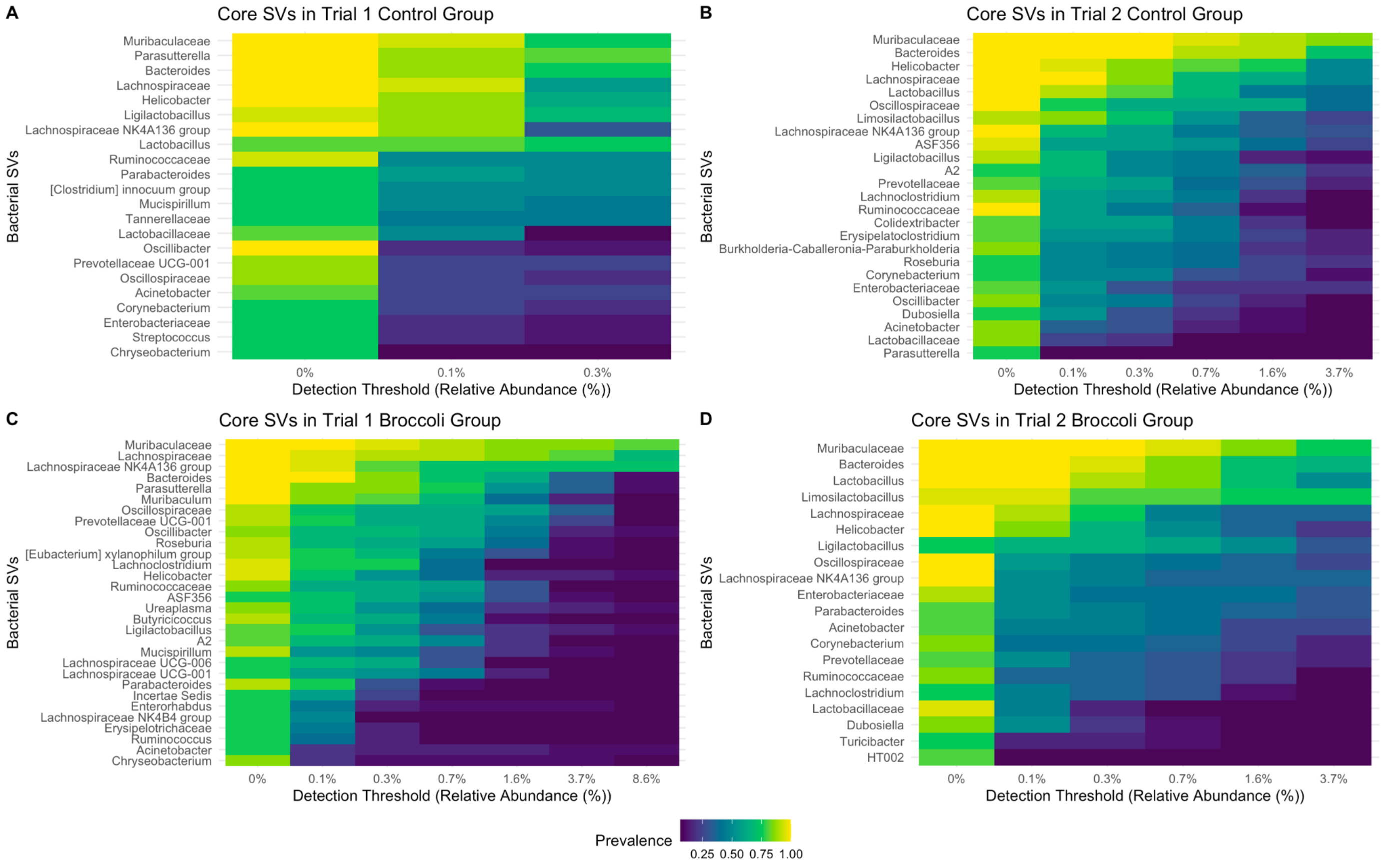


**Figure S5. Core bacterial sequence variants (SV) within the gastrointestinal tracts of 4 week-old (trial 1) or 7-week-old (trial 2) IL-10-ko mice fed control diets or broccoli sprout diets in a Crohn's model of inflammation.** Detection threshold = 1/1000; prevalence threshold = 70/100. Graphics made using microbiome, phyloseq, dplyr, and ggplot2 packages in R.

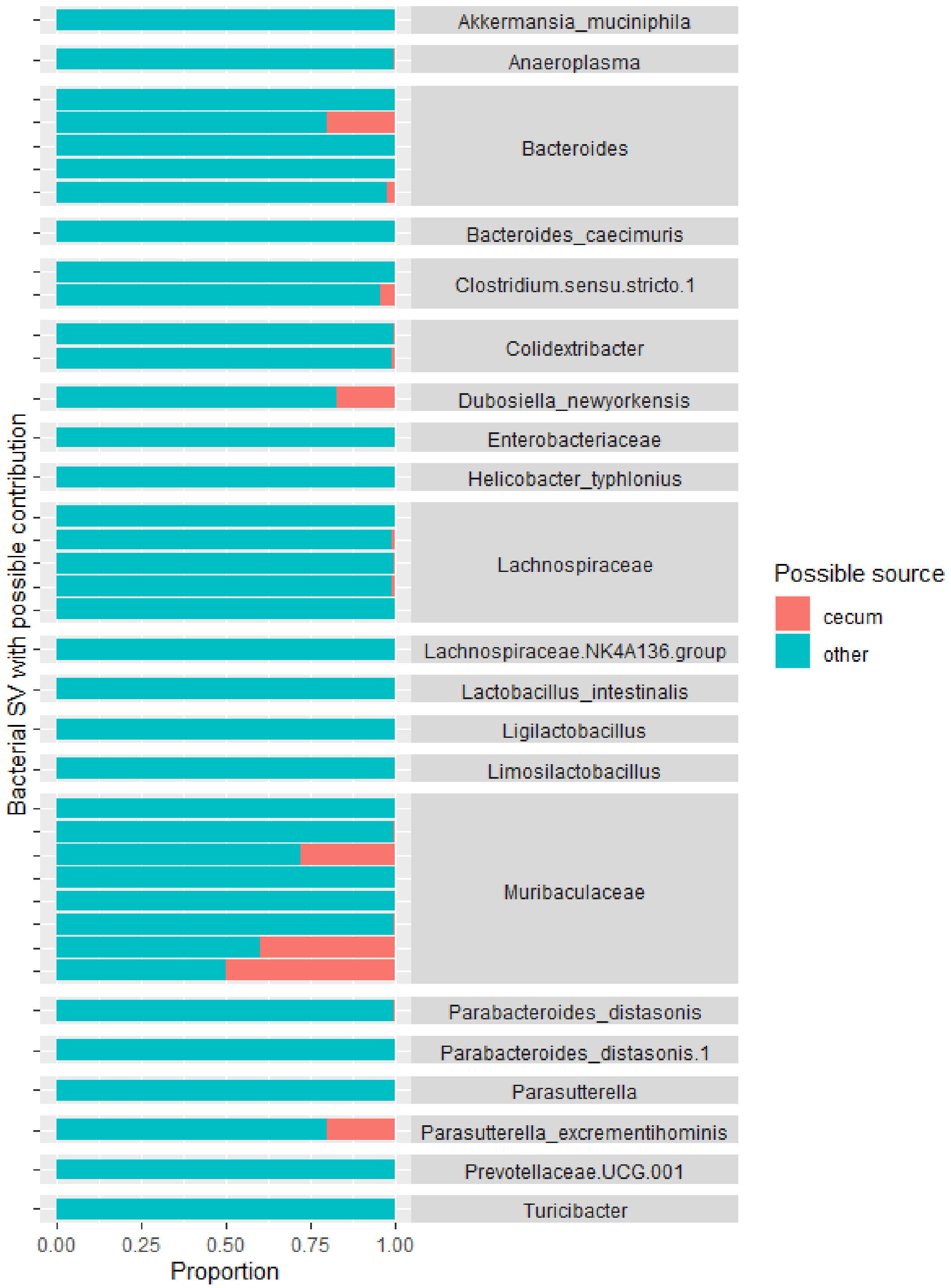


**Figure S6. Bacterial sequence variants which are hypothesized to be sourced from the cecum and creating population sinks in other locations in the intestines of mice.** Potential source SVs were identified with the SourceTracker algorithm modified for the R platform. A total of 38 SVs were identified. Two treatment groups were used in a 3-week immunological model of acute colitis: control diet, and control diet adjusted with 10% by weight raw broccoli sprouts. Bacterial communities were sampled from several locations in the gastrointestinal tract at the end of the study.

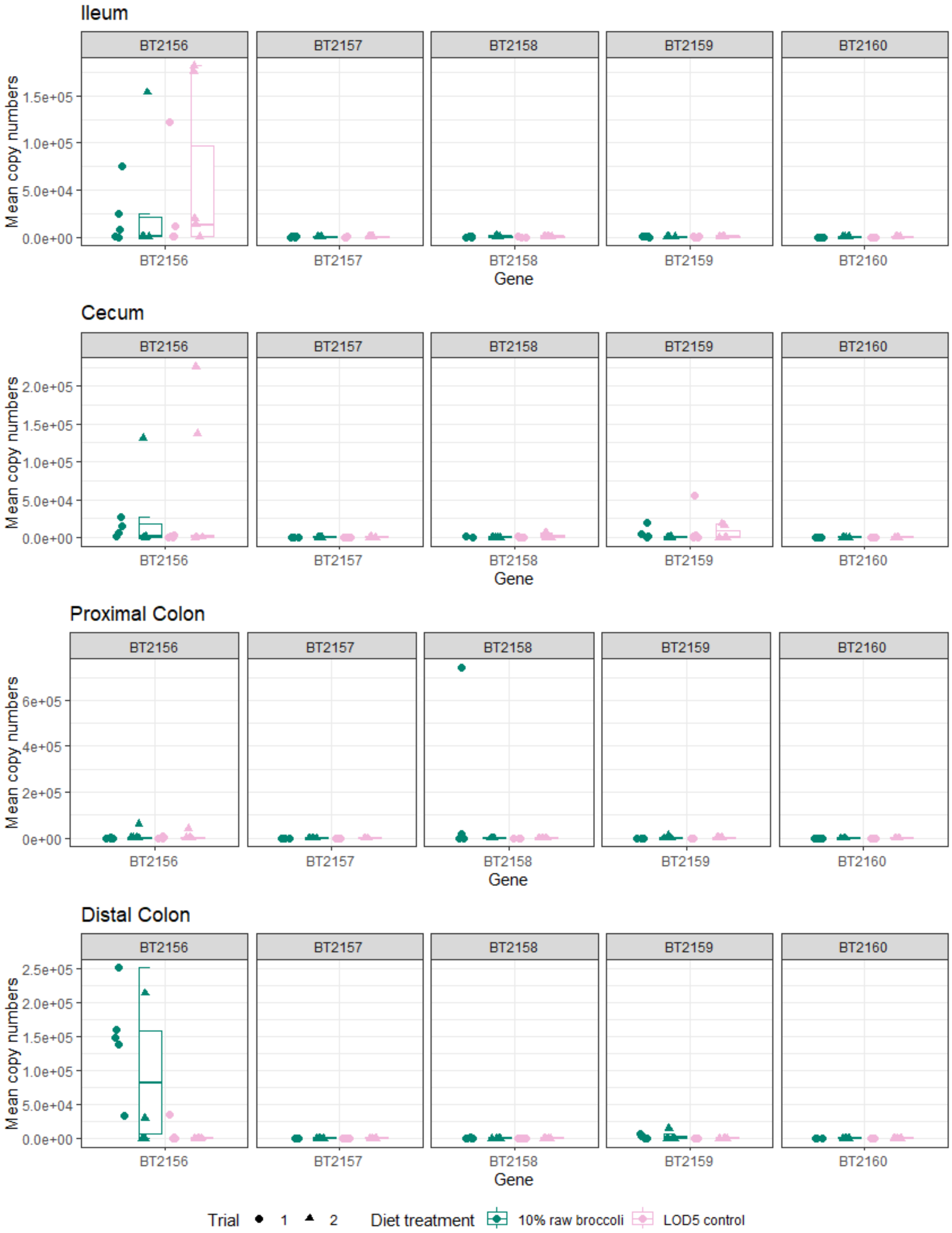


**Figure S7: qPCR results for the Bacterial operon BT2159-BT2156 for *Bacteroides thetaiotaomicron* (VPI-5482) in within a specific anatomical location along the gastrointestinal tract of IL-10-KO mice fed control diets or broccoli sprout diets beginning at 4 or 7 weeks of age.** Note, mean copy number scales vary between charts.

A

B

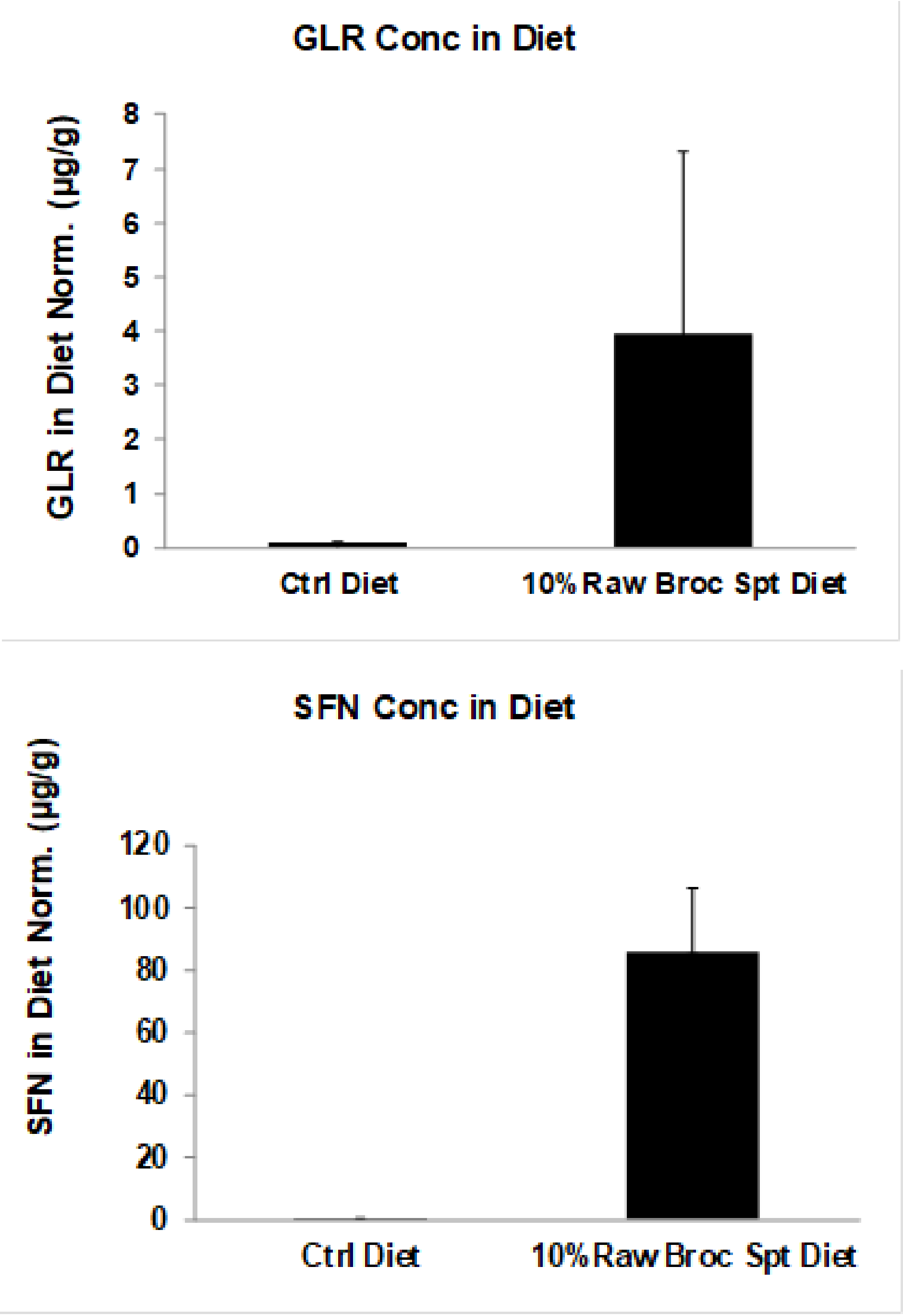


**Figure S8.** Glucoraphanin (GLR) and sulforaphane (SFN) concentration in the 5LOD control diet and the 10% (w/w) raw broccoli sprout diet, as measured by LC/MS.

**Table S1 Targets for quantitative polymerase chain reaction (qPCR) of glucosinolate metabolizing enzymes in *Bacteroides thetaiotaomicron*.**

| Set name | Forward Primer | Reverse Primer | Amplicon size, gene | Reference |
|---|---|---|---|---|
| BT2160 | BT2160_fwd: CAGAATGCCA CACGCGAAAC | BT2160_rev: TGAATGCCGG AGGAAACGA C | 127 bp, SusR4, Transcriptional regulator protein on inner membrane. Controls genes BT2159-BT2156. | (Liou et al. 2020) |
| genebloc k | TGCTGATATTCAGAATGCCACACGCGAAACAGCCTCCCTGCAGGCACTTGCACTG ATACAATATGAGGAA AATAATTTAGCGGATGCTTTCAAATTTACTCAGTCGGCTATTGATGATGTCGTTTC CTCCGGCATTCATT TCCGGGCA | | | |
| Protocol | 1) 1 cycle at 50°C for 2 min,<br>2) 1 cycle at 95°C for 1 min,<br>3) 40 cycles at 95°C for 15 s, 60°C for 30 s and 72°C for 30 s, followed by a plate read,<br>4) melt curve | | | |

| Set name | Forward Primer | Reverse Primer | Amplicon size, gene | Reference |
|---|---|---|---|---|
| BT2159 | BT2159_ fwd TGCGATACA GATCCTACC ACGC | BT2159_rev CAATGTGAAGA GCCCGACAACC | 123 bp, Nicotinamide-dependent oxidoreductase. Cytoplasmic. | (Liou et al. 2020) |
| genebloc k | AGATTGCCTATATTTGCGATACAGATCCTACCAC GCGCGAACTTGCTAAAAAAATATCTCCTGAATCTCTTGTAGTAGAGAATGATCAA AAAATCTTTGAAGAC GAGAGTGTACAGGTTGTCGGGCTCTTCACATTGGCAGACTCAAG | | | |
| Protocol | 1) 1 cycle at 50°C for 2 min,<br>2) 1 cycle at 95°C for 1 min,<br>3) 40 cycles at 95°C for 15 s, 60°C for 30 s and 72°C for 30 s, followed by a plate read,<br>4) melt curve | | | |

| Set name | Forward Primer | Reverse Primer | Amplicon size, gene | Reference |
|---|---|---|---|---|
| BT2158 | BT2158_fwd CGAAACAAT TTGCAGCCG AAC | BT2158_rev GGCAACTTCCA TCCTTCACG | 58 bp, Nicotinamide-dependent oxidoreductase. Periplasmic. | (Liou et al. 2020) |
| genebloc k | TTCATGATGGTCACCCGTCATTCAATAAGACCTGGACAGATCCGATCAA CGCGAAACAATTTGCAGCCGAACTGGTGAAACATAATTATCGTGAAGGATGGAA GTTGCCTGATATGCCACGATAA | | | |
| Protocol | 1) 1 cycle at 50°C for 2 min,<br>2) 1 cycle at 95°C for 1 min,<br>3) 40 cycles at 95°C for 15 s, 60°C for 30 s and 72°C for 20 s, followed by a plate read, | | | |

| | |
|---|---|
| | 4) melt curve |

| Set name | Forward Primer | Reverse Primer | Amplicon size, gene | Reference |
|---|---|---|---|---|
| BT2157 | BT2157_fwd TGCAAGCCAGC AAATTCAGC | BT2157_rev CAGTCCAGAAC TTTCACGCG | 151 bp, Glycosyl hydrolase. Outer membrane lipoprotein. | (Liou et al. 2020) |
| genebloc k | ACCTTTTGCAAGCCAGCAAATTCAGCCAGGACAAATGGCCGTTGGCTTTCGAACT GCTGAATAATTGCGGTGGCGAAAACCACGAAGGATTTATCGGAATGCAGGATCA CGGTGATGACGTTTGG TTCCGCAATATCCGCGTGAAAGTTCTGGACTGA | | | |
| Protocol | 1) 1 cycle at 50°C for 2 min,<br>2) 1 cycle at 95°C for 1 min,<br>3) 40 cycles at 95°C for 15 s, 60°C for 30 s and 72°C for 30 s, followed by a plate read,<br>4) melt curve | | | |

| Set name | Forward Primer | Reverse Primer | Amplicon size, gene | Reference |
|---|---|---|---|---|
| BT2156 | BT2156_fwd CTGCCGGGCTGA AGGTTTTATC | BT2156_rev TCAGCAATACAC TGGTCCCACC | 112 bp, Sugar phosphate isomerase. | (Liou et al. 2020) |
| geneblock | TGTTGAATCTGCCGGGCTGAAGGTTTTATCCTCACATTGCACAAGAGGATTGTCGA AAGAAGAATT AGCTTCCGGTGATTTTTCAAGTTCACTTCAATGGTGGGACCAGTGTATTGCTGATC ATA | | | |
| Protocol | 1) 1 cycle at 50°C for 2 min,<br>2) 1 cycle at 95°C for 1 min,<br>3) 40 cycles at 95°C for 15 s, 60°C for 30 s and 72°C for 25 s, followed by a plate read,<br>4) melt curve | | | |
